# Regulation of skeletal muscle metabolism and contraction performance via teneurin-latrophilin action

**DOI:** 10.1101/2021.10.25.465698

**Authors:** Andrea L. Reid, David W. Hogg, Thomas L. Dodsworth, Yani Chen, Ross M. Reid, Mei Xu, Mia Husic, Peggy R. Biga, Andrew Slee, Leslie T. Buck, Dalia Barsyte-Lovejoy, Marius Locke, David A. Lovejoy

**Affiliations:** Department of Cell and Systems Biology, University of Toronto, Toronto, ON Canada; Department of Biology, University of Alabama, Birmingham AL USA; Protagenic Therapeutics, Inc, Lexington MA USA; Structural Genomics Consortium, University of Toronto, Toronto, ON Canada; Department of Kinesiology, University of Toronto, Toronto, ON Canada.

**Keywords:** TCAP, GPCR, peptides, diabetes, calcium, CRISPR, siRNA, mitochondria

## Abstract

Skeletal muscle regulation is responsible for voluntary muscular movement in vertebrates. The genes of two essential proteins, teneurins and latrophilins (LPHN), evolving in ancestors of multicellular animals, form a ligand-receptor pair, and are now shown to be required for skeletal muscle function. Teneurins possess a bioactive peptide, termed the teneurin C-terminal associated peptide (TCAP) that interacts with the LPHNs to regulate skeletal muscle contractility strength and fatigue by an insulin-independent glucose importation mechanism. CRISPR-based knockouts and siRNA-associated knockdowns of LPHN-1 and-3 shows that TCAP stimulates an LPHN-mediated cytosolic Ca^2+^ signal transduction cascade to increase energy metabolism and enhance skeletal muscle function via increases in type-1 oxidative fiber formation and reduce the fatigue response. Thus, the teneurin/TCAP-LPHN system is presented as a novel mechanism likely to regulate the energy requirements and performance of skeletal muscle.

## Introduction

Skeletal muscle is critical for all voluntary behaviours and is derived from the earliest contractile proteins present in the ancestral single-celled heterotrophs. Enhanced contractile strength and efficient energy metabolism among these primitive skeletal muscle cells were critical for both locomotion and feeding [1,2]. Because of these integrated requirements for the evolutionary success of early metazoans, we have postulated that essential intercellular signaling systems originating phylogenetically early, conferred a selective advantage upon these basal heterotrophs by linking sensory and motor functions with cell metabolism [3]. Numerous studies have indicated that the teneurins and their receptors, the latrophilins (LPHN) are part of an ancient regulatory system that modulates cell adhesion and metabolism. The introduction of the teneurin and LPHN genes into multicellular animals occurred via lateral gene transfer from prokaryotes into a single-celled ancestor of metazoans [4-9]. Thus, the teneurins and LPHNs were evolutionarily poised to play a seminal role in the development and coordination of cell-to-cell communication, adhesion and metabolic activities.

Teneurins are essential for development and maintenance of the central nervous system (CNS) [10-18]. Comprising a family of four paralogous proteins in vertebrates, the teneurins possess several functional domains that confer specialized actions on their extracellular and intracellular regions [19-21]. As type-II proteins, their carboxyl terminus is displaced extracellularly. The most distal region contains a β-barrel structure unique to metazoans, but is similar to that found in prokaryotic Tc-toxins [9,14,22-25]. Associated with this structure lies an extended amino-acid chain termed the ‘teneurin C-terminal associated peptide’ (TCAP) [22-24]. The TCAPs possess primary structure similarity to the Secretin superfamily of peptides that include, not only secretin paralogues such as vasoactive intestinal peptide (VIP), growth hormone-releasing hormone (GHRH), glucagon and pituitary adenylate cyclase activating peptide (PACAP), but also the calcitonin and corticotropin-releasing factor (CRF) families. One of the distinguishing aspects of this peptide superfamily are their roles in the coordination of sensory, motor and energy metabolism [3,26,27].

The LPHNs are G-protein-coupled receptors (GPCR) belonging to the Adhesion GPCR family (ADGRL) and are cognate receptors of the teneurins in vertebrates [28-30]. LPHNs have three distinct paralogous forms (LPHN1-3) and can bind to the C-terminal region of the teneurins, which include the TCAP region. For example, Teneurin-2 and LPHN-1 binds with nanomolar affinity at the lectin-domain of LPHN-1[31]. A splice variant of C-terminal domain of teneurin-2, also termed ‘LPHN1-associated synaptic surface organizer’ (Lasso), binds to LPHN with, likewise, high affinity in neurons. Moreover, transgenic over-expression of both TCAP-1 and the hormone-binding domain (HBD) of LPHN-1 results in co-precipitation of both transgenic proteins indicating an interaction between TCAP-1 and LPHN-1 [32]. Recent structural studies of the teneurins indicate that the TCAP region may be auto-catalytically cleaved from the teneurins after interaction with the LPHNs [14,22], or could be the result of a distinct teneurin splice variant resulting in the mRNA expression of the terminal exon that includes the TCAP sequence [33,34].

Regardless of the mechanism of TCAP release, the expected TCAP mature peptide, based on its genomic sequence, has distinct biological properties. Synthetic TCAP-1 regulates cytoskeletal elements and energy metabolism in neurons critical for neuroplastic modulation in the central nervous system (CNS). In rats, TCAP-1 modifies dendritic arborization and spine density in the hippocampus [35,36], findings that were corroborated in primary cultures of rat embryonic hippocampal tissues that exhibited increased filopodia, neurite and axon development [33,37,38]. Thus, these actions indicate a role of TCAP-1 in CNS energy metabolism. Moreover, subcutaneous administration of TCAP-1 into rats increases brain glucose uptake as assessed by functional positron emission tomography (fPET). These observations were corroborated by the expected decreased serum glucose and insulin levels in rats, and in cell culture studies showing that TCAP-1 stimulates glucose uptake by increased glucose transporter transit to the membrane and subsequently increases in ATP turnover providing increased energy for the neurons [39].

However, given the evolutionary history of the teneurins, it is plausible that the teneurins, LPHNs, and TCAP could also play a role in the regulation of skeletal muscle. Skeletal muscle is one of the most important sites of glucose metabolism and is responsible for 40% of glucose-associated energy requirements [39] and 80% of glucose uptake under insulin-stimulated conditions [40]. Muscle function and metabolism are intrinsically linked, as evidenced by metabolic syndromes that result in poor muscle function and degradation. A key example of this is demonstrated in patients with type-II diabetes where patients have reduced skeletal muscle function in the grip force test compared to non-diabetic patients [41,42].

Based on these previous findings, we investigated the role of TCAP-1 on skeletal muscle function for the first time. We demonstrate that skeletal muscle possesses the critical elements of teneurin-LPHN interaction, and show that TCAP-1 regulates skeletal muscle contractile kinetics *in vivo* in rats. These studies are supplemented by *in vitro* studies, using the mouse skeletal cell line, C2C12, to show that TCAP regulates intracellular skeletal Ca^2+^ flux similar to that shown in neurons [3,39]. Moreover, like neurons, the TCAP-mediated Ca^+2^ response leads to increased glucose metabolism and mitochondrial activation, but results in skeletal muscle fiber regulation. We posit that the teneurin-LPHN interaction is essential for skeletal muscle physiology and regulates skeletal muscle performance.

## Materials and Methods

### Peptide Synthesis and Solubilization

Both peptides; rat/mouse TCAP-1 and scrambled TCAP-1 (Fig. 1), were synthesized commercially by AmbioPharm, Inc. and prepared as an acetylated salt at 95% purity. Peptides were solubilized in saline after alkalization as previously described [37] then diluted into the required media for *in vitro* or *in vivo* studies (see below). The primary structure of all four rat and mouse TCAPs are identical to each other (Fig.1A) and possess a 73-83% sequence identity among the overall sequences, although most of these changes reflect homologous and conservative substitutions. For this reason, TCAP-1 was used in both rat and mouse preparations. Synthetic rat/mouse TCAP was prepared with an initial N-terminal pyroglutamic acid to inhibit N-terminal-directed peptidases, and a C-terminal amidated-residue as expected based on the genomic sequence [24,37]. As a control peptide, we have utilized a scrambled (sc) amino acid sequence version of rat/mouse TCAP-1 where each residue, with the exception of the initial pyroglutamyl residue (pE), was randomized in its placement within the peptide (Fig. 1B). This sc-TCAP-1 has been used in previous studies to establish an additional level of controls to ensure that TCAP-1 is not affecting non-specific (e.g. oligopeptide transporters; non-target receptors) actions. The vehicle included sc-TCAP solubilized in 0.9% saline or cell culture medium, unless otherwise stated.

**Figure 1.**
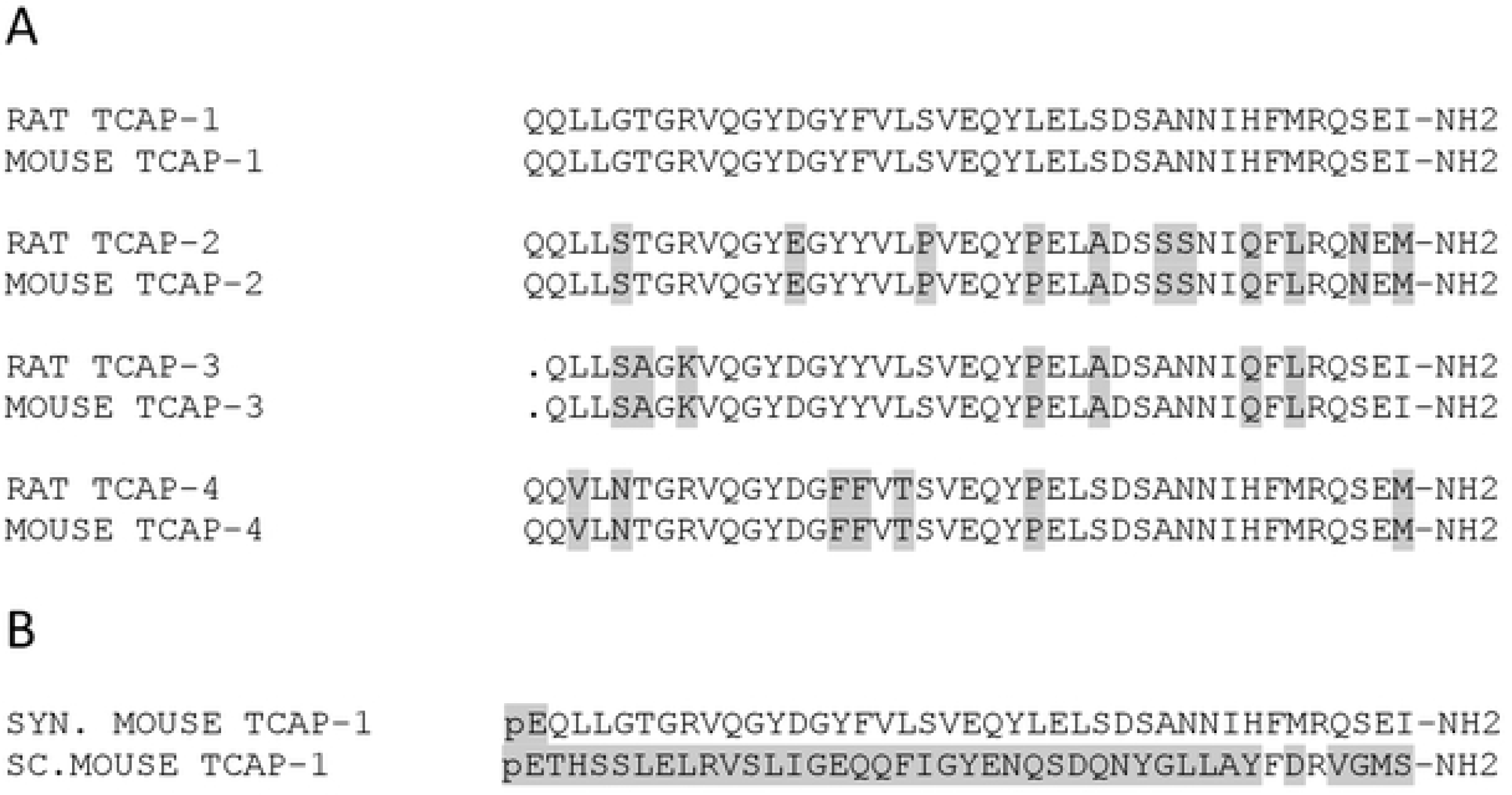
Primary structures of rat and mouse TCAP peptides. **A**. Comparison of the amino acid sequences of mouse and rat TCAPs. **B**. Primary structure of the peptides used in this study. Grey boxed regions indicates regions of identity relative to the rat/mouse TCAP isoforms.

### Animals

Male adult Sprague-Dawley (SD) rats (Charles River, Canada) were used for the short-term and long-term muscle function studies. The metabolic and endocrine studies of TCAP-1 on rats were approved by the University of Toronto Animal Care Committee (UACC) under the auspices of the Canadian Council of Animal Care. Male adult Wistar rats (∼250 g) (Charles River, USA) were used for the functional positron emission tomography (fPET) studies performed by Molecular Imaging, Inc. (Ann Arbor, MI, USA) and approved by the American Association of Animal Laboratory Care (Hogg et al., 2019). In both sets of studies, animals were weighed weekly and monitored for any signs of distress or illness (e.g. loss of hair, extreme weight loss, abnormal behaviours). However, no animals showed overt indications of stress and all were utilized for these studies.

### In Vivo Studies

The short-term application of TCAP-1 utilized 16 male adult SD rats (250g) that were acclimated for 1 week (w) on a 12:12 light-dark (LD) cycle. For 5 days (d) daily, the animals were treated with either vehicle or TCAP-1 (10 nmoles/kg) by subcutaneous (SC) injection in the intrascapular region. Animals were tested for muscle function by electrical muscle stimulation (see below) 3d after the last treatment. Animals were immediately euthanized afterward. For the long-term study, 20 adult male SD rats (350g), acclimated for 1w on a 12:12 LD cycle and were treated with either vehicle or intrascapular TCAP-1 (SC; 25 nmoles/kg) for 3 months (1 injection/week). Muscle function by electrical muscle stimulation was tested 2w after the last treatment. Animals were immediately euthanized after electrical muscle stimulation studies (see below).

### PCR Expression of teneurin and LPHN

RNA was extracted from tibialis anterior (TA) muscles using TRIzol (Thermo Scientific, Waltham, MA, USA) following the manufacturer’s instructions. The PCR reaction mix included 5 μL cDNA, 2 μL forward primer and 2 μL reverse primer (Invitrogen; Table 1), 14.2 μL water (Sigma, Oakville, ON), 3 μL 10x Taq Buffer with KCl (Thermo Scientific), 1.8 μL MgCl_2_ (Thermo Scientific), 1 μL deoxynucleotide Solution Mix and 0.5 μL Taq DNA Polymerase (New England Biolabs). The reactions were incubated in an Eppendorf Mastercycler Gradient Thermal Cycler for 7m at 95°C; followed by 35 cycles of 60s at 95°C, 90s at 67°C, and 35s at 72°C; and then held at 4°C. cDNA samples were resolved on a 3% agarose gel at 100 V for 1.5h and visualized using a Bio-Rad ChemiDoc MP System with 0.5s exposure.

**Table 1.**
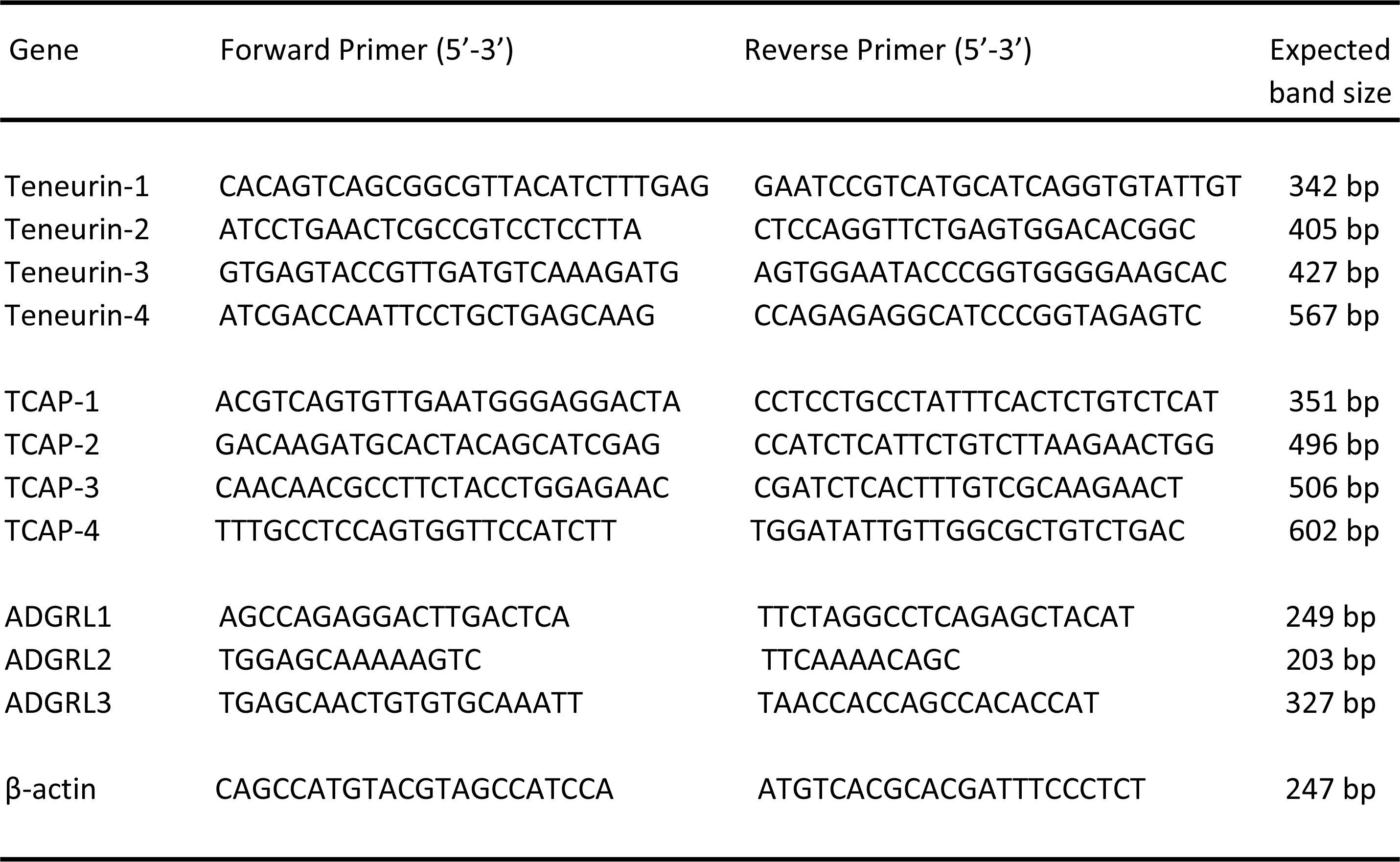
Primers used for in vitro and in vitro RT-PCR analyses. Forward and reverse primer pairs for the four teneurins, TCAPs, three ADGRLs, and β-actin control are indicated.

### Histological Studies

TA muscle was excised then flash-frozen in liquid nitrogen cooled-isopentane. The tissue was cryo-sectioned at 10 μm at -20°C and transferred to coverslides and fixed using ice-cold methanol. Following blocking for 1h with 10% normal goat serum (NGS: Cell Signaling, Inc.), the primary antibody (Table 2), diluted in 1% NGS, was added and incubated overnight (ON) at 4°C. Subsequently, after 3 phosphate-buffered saline (PBS) washes, the secondary antibody was added and incubated for 1h at room temperature (RT) in the dark. Coverslips were attached, and the tissue imaged using confocal microscopy (Leica TCS-SP8) at 400x magnification. For fluorescence analyzes of protein expression, Image J software was used to measure arbitrary fluorescent units (AFU), where changes of AFU are proportional to protein expression changes. A total of 5 slides were quantified with 8 regions of interest (ROI) investigated. ROI was defined as regions with multiple cell interactions free of artifacts.

**Table 2.** Primary and secondary antibodies used in western blot analyzes.

| Name | Primary Antibody | Secondary Antibody | Expected band size |
| --- | --- | --- | --- |
| LPHN-1 | Goat polyclonal IgG | Donkey anti-goat HRP-linked | 130 kDa |
| LPHN-2 | Goat polyclonal IgG | Donkey anti-goat HRP-linked | 163 kDa |
| LPHN-3 | Rabbit polyclonal IgG | Donkey anti-rabbit HRP-linked | 162 kDa |
| SDH-A | Mouse monoclonal IgG | Rabbit anti-mouse HRP linked | 70 kDa |
| β-actin | Mouse monoclonal IgG | Rabbit anti-mouse HRP linked | 42kDa |

### Functional Positron Emission Tomography (fPET)

Male Wistar rats were treated with either vehicle or TCAP-1 (10 nmoles/kg) by intrascapular SC injection. 3d after treatment, 1 mCi of [^18^F]-2-deoxyglucose (^18^F-DG) radiotracer (IBA Molecular) was administered intravenously (IV) under anesthesia. fPET scans were performed using a Siemens Inveon microPET small animal PET scanner using the protocol as previously described [39]. Briefly, body temperature was maintained with a thermostat-regulated recirculating water-heated pad. Static emission data was acquired for 20m. The PET list mode data was converted to 2-dimensional (2D) sinograms, corrected for random coincidences, and normalized for scanner uniformity. The PET image analysis was performed using the Amira 5.5.0 analysis software package. For whole body ROIs (regions of high ^18^F-DG uptake), a low threshold was set to delineate specific signals in the whole body while eliminating background. The total PET counts were calculated from all voxels within the segmented volumes of interest. These images were then compiled into 3-dimensional (3D) projections, thus allowing for accurate analyzes of muscle tissue. Fluorescence of the mean pixel was calibrated to volume of muscle being analyzed (mean pixel fluorescence/mm^3^).

### NADH staining and analysis

TA muscles from the treated SD rats were cryo-sectioned at 10 μm thickness as described above. Cryo-sections were washed 2x with PBS, then 0.2% NBT solution in PBS (containing 0.1% NADH; Sigma, Oakville, ON) was added and incubated for 30m at 37°C. Slides were washed 2x in PBS before mounted, imaged at 100x magnification and analyzed on Image J software for pixel density, where darker pixels represent higher levels of NADH. Expression of NADH was analyzed based on a minimum of 100 fibres per tissue, with a minimum of 3-4 tissues analyzed for each group.

### Muscle function and integrity testing by electrical muscle stimulation

The electrical muscle stimulation protocol was followed as described by Holwerda and Locke [43] with minor modifications. Briefly, animals were anesthetized with 5% isofluorane in 1L/min O_2_, and subsequently positioned into the testing apparatus. A 25g needle was inserted through the soft tissue of the knee in order to ensure a stable position. The foot was placed on a lever attached to a servomotor and taped in position. Electrodes were placed below the skin but adjacent to the TA muscle. Dynamic Muscle Control (DMC; Version 5; Aurora Scientific) software was used for electrical stimulation and analyzes. The correct voltages for peak tetanic tension was established by increasing voltage by 1V increments until optimal tetanus twitch was achieved. The test began with a single tetanus and single twitch protocol to establish the baseline followed by a 6-min fatigue protocol (8V, 200 Hz, 300 ms). After the termination of the protocol, tetanic and twitch tensions were recorded at 0, 1, and 5 mins. Animals were immediately euthanized after recovery measurements were recorded.

### Quantitative Reverse-Transcription Polymerase Chain Reaction (qRT-PCR)

TA muscle mRNA and cDNA were prepared as previously described (see above). The cDNA from all samples was used to prepare pools to establish standard curves of each gene. The cDNA pool or cDNA samples were mixed with MasterMix containing SYBR select. The reactions were loaded in a 384-well PCR plate and run in a BioRAD qRT thermal cycler for 2m at 50°C, 7m at 95°C; followed by 39 cycles of 60s at 95°C, 90s at 67°C, and 35s at 72°C. Melting curves were established by a step-wise gradient from 60-90°C. The myosin heavy chain (MHC) isoforms, MHCI, MHCIIa, MHCIIx and MHCIIb were analyzed by real-time PCR using the mRNA and cDNA prepared above (see Table 3).

**Table 3.**
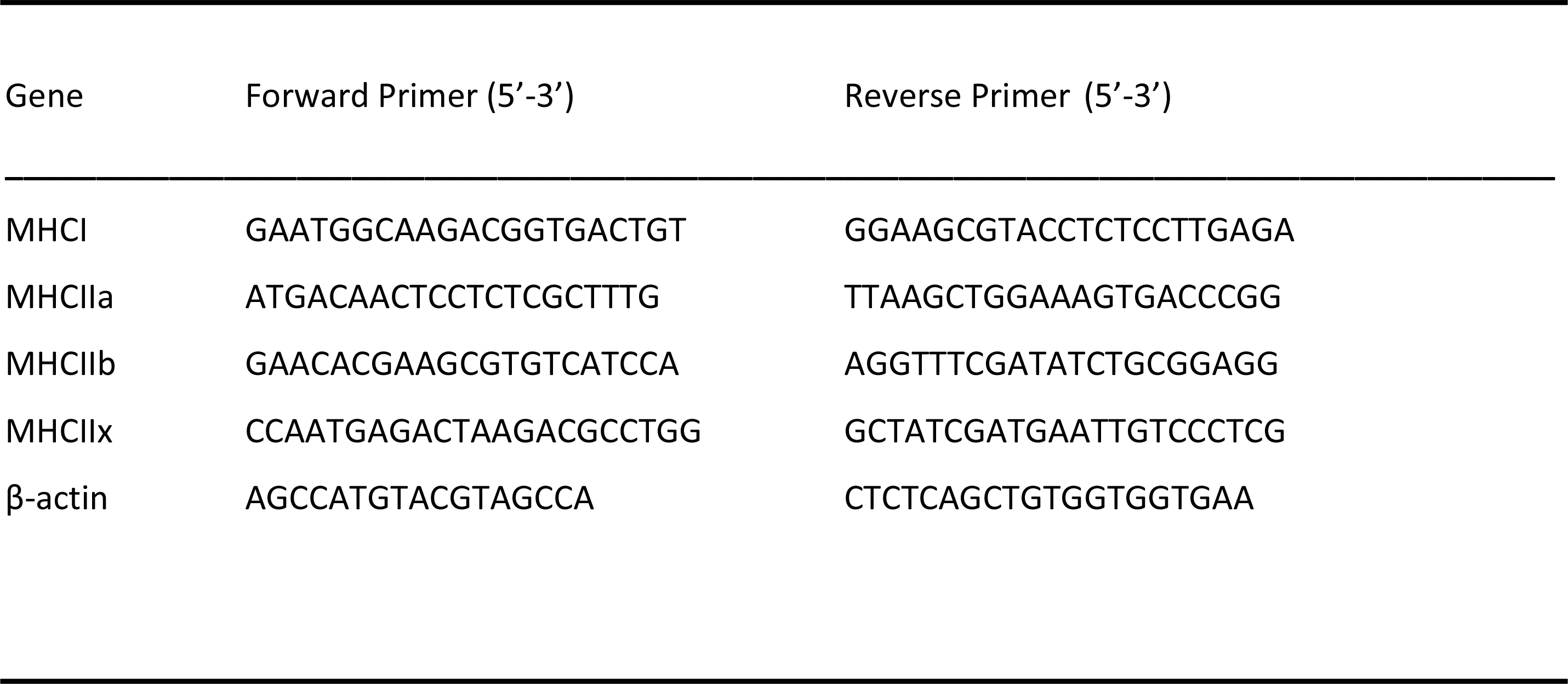
Rat MHC isoforms and β-actin control primers used for qRT-PCR. Forward and reverse primer pairs are indicated for the four MHC isoforms.

### In Vitro Studies

#### Culture and cDNA analyses of C2C12 cell line

The immortalized murine skeletal muscle cell line, C2C12, was used for all *in vitro* studies. Cells were maintained at 60-70% confluency with Dulbecco’s Modified Eagle Medium (DMEM) supplemented with 20% fetal bovine serum (FBS) and a penicillin/streptomycin antibiotic combination (Invitrogen, Burlington, ON, Canada). To induce differentiation, the media was changed to DMEM supplemented with 10% horse serum (HS) with the penicillin/streptomycin antibiotic combination, and the cells were allowed to differentiate for 6d (media replaced every 24 hrs). For treatment, cells were serum-starved for 3h and then treated with either vehicle or TCAP-1 (100 nM). The identification of teneurin, TCAP and LPHN cDNAs in C2C12 cells was performed with mRNA extracted from differentiated mouse C2C12 cells using the method described above using the primer sequences indicated in Table 1.

#### Live-cell calcium imaging in C2C12 myotubules

The C2C12 skeletal cells were grown and differentiated on poly-D-lysine-coated 25 mm round No. 1 glass coverslips (Warner Instruments, Hamden, CT, USA). Changes in intracellular Ca^2+^ were assessed using the membrane-permeable fluorescent indicator fluo-4, AM (Invitrogen, Burlington, ON, Canada). Cells were loaded with fluo-4 by incubating coverslips in DMEM containing 10 μM fluo-4 for 30m (37°C) followed by a 15m wash in Locke’s Buffer (305-310 mOsmol at 22°C). In experiments assessing changes in intracellular Ca^2+^, coverslips were placed in a flow-through bath chamber (RC-40HP, Warner Instruments, Hamden, CT, USA) of an inverted microscope (Axio Observer Z1, Zeiss, Toronto, ON, Canada) equipped with a 40× oil immersion objective. Cells were continuously bulk-perfused with Locke’s buffer via a gravity drip perfusion system at a rate of 2–3 ml/min at RT. Changes in fluo-4 fluorescence were imaged using a green fluorescent protein (GFP) filter set (Semrock, Rochester, NY, USA) and a X-Cite 120 fluorescence illumination system (Excelitas Technologies, Mississauga, ON, Canada), controlled by Volocity 4.0 imaging software (Quorum Technologies Inc., Guelph, ON, Canada). Fluorescence emissions were detected with an Orca-ER Hamamatsu B/W CCD digital camera (Hamamatsu, Middlesex, NJ, USA). Fluo-4 was excited with a wavelength of 480 nm for 100ms every 3-5s and fluorescence emission was measured at wavelength of 516 nm.

#### Caffeine stimulation experiments

Caffeine action on the cells was used to establish that the cells were viable. Caffeine (4 mM; Sigma-Aldrich, Oakville, ON) was applied to C2C12 myotubes to stimulate Ca^2+^ release from the sarcoplasmic reticulum (SR). Cells were either pre-treated with TCAP-1 (100 nM) or vehicle for 1h before stimulation with caffeine. Using Velocity 4.0 imaging software, ROIs were taken from cytosolic regions within the myotubules (n= 4 coverslips, 4-5 ROIs per coverslip).

#### GLUT4 Immunocytochemical studies

C2C12 cells were differentiated, as described above. After 3h of serum starvation, myotubules were treated either with vehicle, TCAP-1 (100 nM) or insulin (100 nM) for 15 or 30m. Cells were then fixed using 4% paraformaldehyde and subsequently blocked with 10% NGS for 1h at RT. The GLUT4 primary antibody, diluted in 1% NGS, was added to the cells and incubated at 4°C overnight (OT). Following 4 PBS washes, the secondary antibody (diluted in 1% NGS) was added and incubated 1h at RT. The coverslips were mounted using DAPI-containing Vectashield. Slides were imaged on a confocal microscope with a 40x oil objective. The images were analyzed using Image J, where myotubules were selected as ROIs and were analyzed for red pixel intensity values, representing GLUT4 levels, and normalized to area size (n= 3-4 coverslips per treatment, 7-8 myotubules per coverslip). For the IP3R inhibitor, 2-aminoethoxydiphenyl borate (2-APB; Sigma Aldrich, Oakville, ON) experiments, 2-APB (100 µM) was applied for 4m before the start of treatment with either sham (Locke’s buffer) or TCAP-1 (100 nM), containing 2-APB for continuous blocking of IP3R.

#### Radioactive glucose uptake

The ^3^H-2-deoxyglucose uptake protocol was followed as previously described with minor modifications [44,45]. At day-6 post-plating, C2C12 myotubules were washed 2x with Locke’s without serum and glucose. The culture was incubated in the Locke’s buffer for 1h at 37°C followed by exposure to 100 nM insulin, 100 nM TCAP-1, 100 nM SC-TCAP-1, or saline. ^3^H-2-deoxyglucose (0.5 μCi/ml) was added to the culture 5m before termination of treatment exposure. Uptake of ^3^H-2-deoxyglucose was stopped immediately after 5m with 3x washes of ice-cold 0.9% saline solution. The cells were digested with 1 mL of 0.05 M NaOH at 0, 30, 45, and 60 min after treatment. Radioactivity of the cell lysates were measured using a beta liquid scintillation counter (Beckman Coulter), and recorded in counts per minute (CPM).

#### Intracellular ATP and NADH assays

ATP assays were conducted using Promega ATP Assay kits (Wisconsin, USA) following the manufacturer’s instructions. Briefly, C2C12 cells were seeded at 10,000 cells/well in 96-well plates. The following day, cells were treated with either vehicle or TCAP-1 (100 nM) and lysed at 0, 15, 30 and 60m after treatment. Ultra-Glo recombinant luciferase (Promega, Wisconsin, USA) was added to the media to determine ATP levels. Fluorescence from blank wells was subtracted from all samples to account for background signal noise. As the fluorescence signal naturally decays over the course of the experiment, TCAP-1-treated cells were compared relatively to the vehicle-treated cells for each time point (n=8). For the resazurin NADH assay, the C2C12 cells were seeded at 10,000 cells /well in 96-well plates. The resazurin assay was started the following day by adding the resazurin solution (525 nM, Sigma) to all wells. Cells were treated with either vehicle or TCAP-1 (100 nM). Fluorescent readings were measured every 5m over 1h, with excitation at 530 nm and emission read at 590 nm. The measurements of blank wells not containing cells were subtracted from all readings.

#### Western blot of succinate dehydrogenase in C2C12 cells

Following TCAP-1 treatments, proteins were extracted from C2C12 cells and lysed with radioimmunoprecipitation assay (RIPA) buffer supplemented with the protease inhibitor, phenylmethylsulfonic fluoride (PMSF) (Cell Signaling Technology, MA USA) and measured using a bicinchonic acid (BCA) protein assay (Pierce BCA Protein Assay; Thermo-Fisher Scientific, Toronto, Canada). All protein extracts were re-suspended in sample buffer and size-fractioned by SDS-PAGE, electro-transferred to nitrocellulose membranes. Afterwards, membranes were incubated with the succinate dehydrogenase (SDH) primary antibody and subsequently treated with secondary antibody conjugated with chemiluminescent tags (Table 2). Following TCAP-1 treatments, C2C12 cells were lysed with 500 μL of RIPA buffer supplemented with PMSF. Cells were harvested and centrifuged at 20,000 rcf for 20m at 4°C. Protein concentrations (as described above) were measured to standardize the dilutions of respective supernatant samples. Samples (15 μg) were re-suspended in sample buffer and size-fractioned by SDS-PAGE (10%) at 100V for 1h. The protein extract was electro-transferred to Hybond-ECL nitrocellulose membranes (Amersham) for 2h at 100V. Membranes were washed 3x with PBS and blocked in 5% milk-PBST (5% w/v non-fat milk powder in PBS with 0.2% Tween20; PBST) at RT for 1h, then incubated with primary antibodies in 1% milk-PBST OT at 4°C. The membranes were washed 3x in PBST at RT and incubated with a horseradish peroxidase (HRP)-conjugated secondary antibody (VWR, Amersham; diluted to 1:7500 in 1% milk-PBST) for 1h at RT then washed 3x in PBST at RT. The proteins were detected by adding chemiluminescence detection reagent (ECL Amersham) to the membranes and exposing onto ECL Hyperfilm (VWR) for 30m.

#### Diacylglycerol (DAG) and inositol triphosphate (IP3) assays

The protocols provided by commercial DAG and IP3 assays (MyBiosource, San Diego, California, USA) were followed. To determine the downstream Ca^2+^ response, 6 replicates of C2C12 cells were prepared using the TCAP-1 treatment protocol described above then treated with either vehicle, the IP3R antagonist, 2-APB, or the phospholipase C inhibitor, U73122. Cell lysates were added to a microELISA plate coated with purified mouse DAG or IP3 antibodies. Subsequently, 3,3’,5,5’-tetramethylbenzidine (TMB) solution was added to detect the HRP-conjugates as colour changes. Finally, sulphuric acid (0.01N) was added to terminate the reaction. The absorbance change was measured at 450 nm by spectrophotometry (SpectraMax Plus, NH, USA). For the IP3R inhibitor, 2-APB (100 µM) was applied before the start of treatment with either sham (Locke’s buffer with scTCAP-1) or TCAP-1 (100 nM). For live-cell fluorescence experiments, C2C12 cells were differentiated and intracellular Ca^2+^ flux was assessed via fluo-4 via a flow-through bath chamber of an inverted microscope. Cells were quantified with a GFP filter set at 480 nm with the fluorescence emission measured at 516 nm.

#### Mitochondrial Ca^2+^ accumulation and membrane potential measurement in C2C12 myotubules

Changes in mitochondrial Ca^2+^ levels were assessed using fluorescent indicator, Rhodamine-2-AM (Rhod-2). C2C12 myotubules were loaded with Rhod-2 by incubating coverslips in DMEM containing 4µM Rhod-2 (from a 1mM stock solution in DMSO with 20% pluronic; Invitrogen-Pluronic™ F-127) for 30m at 22°C. Cells were washed once for 30m at 37°C in Locke’s Buffer Cells then acclimated for 15m at 22°C. To assess changes in mitochondrial Ca^2+^ levels, cells were continuously perfused in a flow-through chamber as indicated previously. Changes in Rhod-2 fluorescence was imaged using a TRITC filter set (Semrock, Rochester, NY, USA) and an X-Cite 120 fluorescence illumination system (Quorum Technologies, Inc. Guelph ON, Canada). Emissions were detected using an Orca-ER Hamamatsu BW CCD digital camera as described above. Rhod-2 was excited at 552nm every 100 ms and measured at 577nm. Multiple ROI were taken from the nuclear regions of the myotubules (n=5, 5-7 ROIs per coverslip). Changes in mitochondrial membrane potential were assessed using R123-based fluorescence. C2C12 myotubules were prepared with by incubating a coverslips in DMEM containing 5 μM R123 for 30m (37°C) followed by a 15m wash in Locke’s Buffer. Changes in R123 fluorescence was imaged using the green GFP filter set using the same experimental configuration as previously described. R123 was excited with a wavelength of 480 nm for 100 ms every 5s and fluorescence emission was measured at 516 nm.

#### SiRNA knockdowns (KD) and CRISPR Knockouts (KO) of LPHN-1 and -3

For the siRNA KD studies, transfection with siRNA oligonucleotides was performed after 4d of C2C12 differentiation using Dharmacon SmartPOOL (Horizon Inc. Canada) siRNA for LPHN1 (L-061299-00-0005), LPHN3 (L-040779-00-0005) and a non-targeting control (D-001810-10-05). Dharmacon SmartPOOL siRNAs targeted against LPHN-1 and-3, glyceraldehyde-3-phosphate (GAPD) and a non-targeting control were re-suspended in 1x siRNA buffer from 20 µM stocks. The stocks were diluted in serum-free and antibiotic-free DMEM to 250 nM. A 7.5µL aliquot of Mirus TranslT-X2 (Mirus Bio LLC) transfection reagent was diluted in 200 µL serum- and antibiotic-free DMEM and incubated at RT for 30m. The mixture was added to C2C12 cells (see above) with a final siRNA concentration of 25 nM. The cells were differentiated in siRNA-containing media for 2d for a total of 6d of differentiation before use in the experiments. For CRISPR studies, single-guided RNA (sgRNA) constructs were designed to target the mouse LPHN-1 and -3 gene at 3 locations (see Fig. 2). C2C12 cells were transfected with sgRNA constructs (Fig. 2C) and a Cas9 plasmid, generating heterogenous pools of transfected cells. The CRISPR/Cas-transfected C2C12 cells (either hetergenous pools or clones) were trypsinized and pelleted for DNA extraction. Genomic DNA was extracted using Lucigen QuickExtract DNA extraction solution (Biosearch Technologies, Inc) according to the manufacturer’s direction. The LPHN-1 and -3 genes were amplified by PCR and digested by T7 endonuclease using the EnGen Mutation Detection Kit (New England Biolabs) according to directions in combination with the custom primers that flank the appropriate CRISPR-targeting regions (Fig. 2C). The fragments were identified as previously described above. Clones that showed low or no WT PCR amplicon were screened for LPHN1 expression by qRT-PCR. Selected clones showing significantly reduced LPHN1 mRNA expression by qRT-PCR and western were termed ‘LPHN1 E5U KO’ and ‘LPHN1 E5D KO’ based on the exon position of mutated site and were used for further study (Fig. 2). The activity of the clones were determined by TCAP-1-induced cytosolic Ca^2+^ flux in the manner described above. Peroxisome proliferator-activated receptor γ coactivator 1α (PGC-1α) expression in C2C12 cells was determined using the qRT-PCR methods described above.

**Figure 2.**
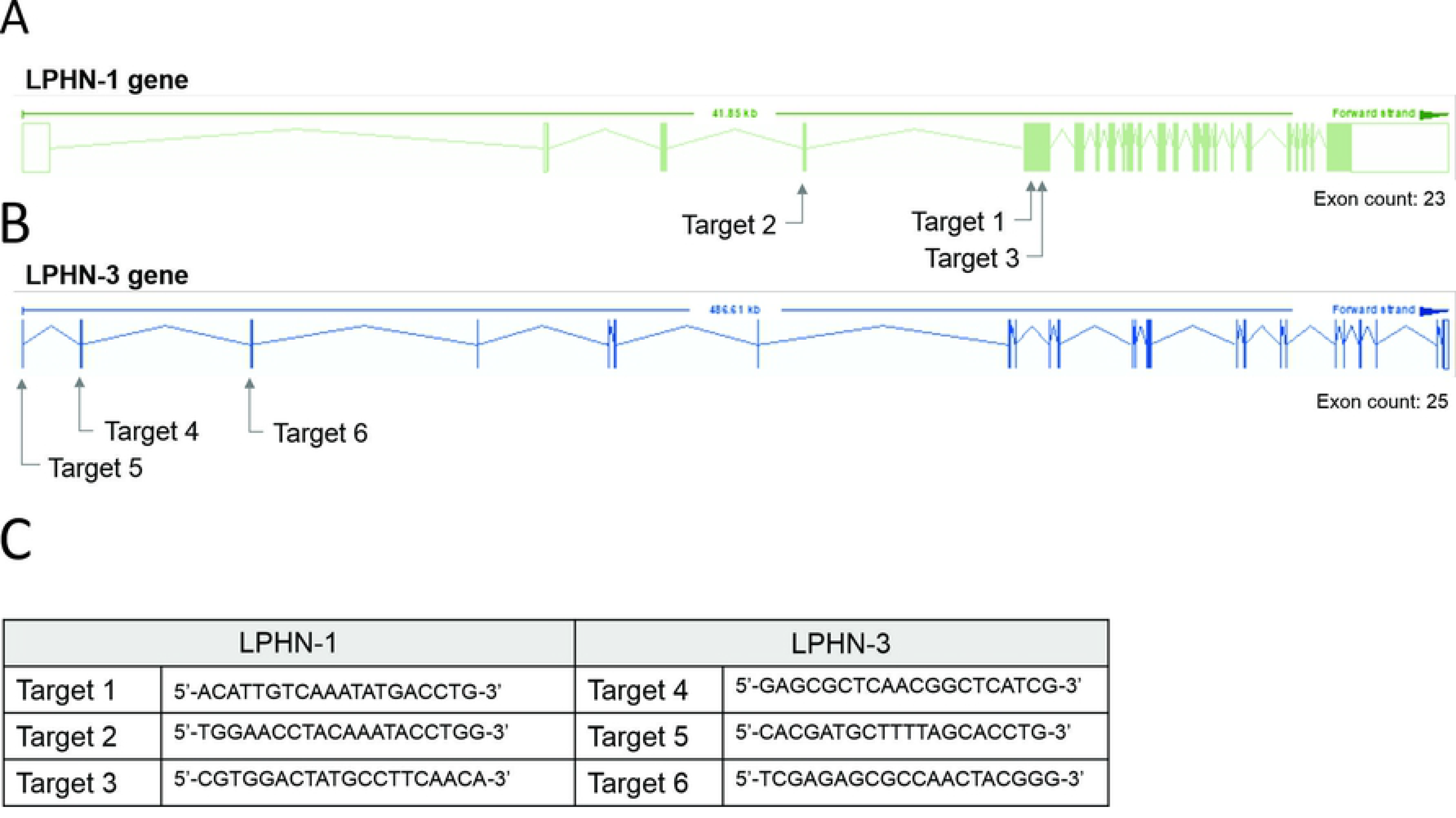
CRISPR-based targets for the mouse LPHN-1 and -3 genomic sequences. **A.** Schematics of LPHN-1 and -3 genomic organization. The oligonucleotides targeting these regions are indicated as arrows. **B.** Sequence of the oligonucleotides used in the CRISPR knockdowns as indicated in ‘A’ above.

### Statistical Analyses

All data on graphs are represented as mean ± SEM. All data were analyzed by Student’s t-test or one or two-way ANOVA, as described within each figure caption. Tukey’s post-hoc test and Sidak’s post-hoc test were used to determine significance in one-way and two-way ANOVA analyzes, respectively. An *a priori* hypothesis of p<0.05 was used as a threshold for statistical significance. GraphPad Prism 7-8 was used to analyze each statistical test.

## Results

The primary structure of rat and mouse TCAP-1 possesses a high degree of homology among the other three paralogues (Fig 1A). Because the primary structure of rat and mouse TCAP-1 is identical, it was used for all studies, as well as a proxy for the other TCAP isoforms. The first studies completed established that the teneurins, TCAP and LPHNs could be expressed in rat skeletal muscle. Secondarily, the physiological role of TCAP-1 in rat skeletal muscle was examined.

### In Vivo Rat Studies

In rat TA muscle mRNA extracts, all 4 teneurin mRNAs were identified based on the PCR primers indicated in Table 1. Teneurins-3 and-4 showed the strongest response, although both teneurins-1 and -2 were present, albiet weakly expressed. In contrast, TCAP-1 and -2 showed a strong signal relative to that indicated by teneurins-1 and -2 whereas TCAP-4 showed a signal consistent with teneurin-4. Although these studies were not quantitative, they do establish that both teneurins and TCAP paralogues are present in rat skeletal muscle. However, importantly, both LPHN-1 and -3 cDNA bands were also strongly expressed, although there was no evidence of LPHN-2 in this preparation (Fig 3A).

**Figure 3.**
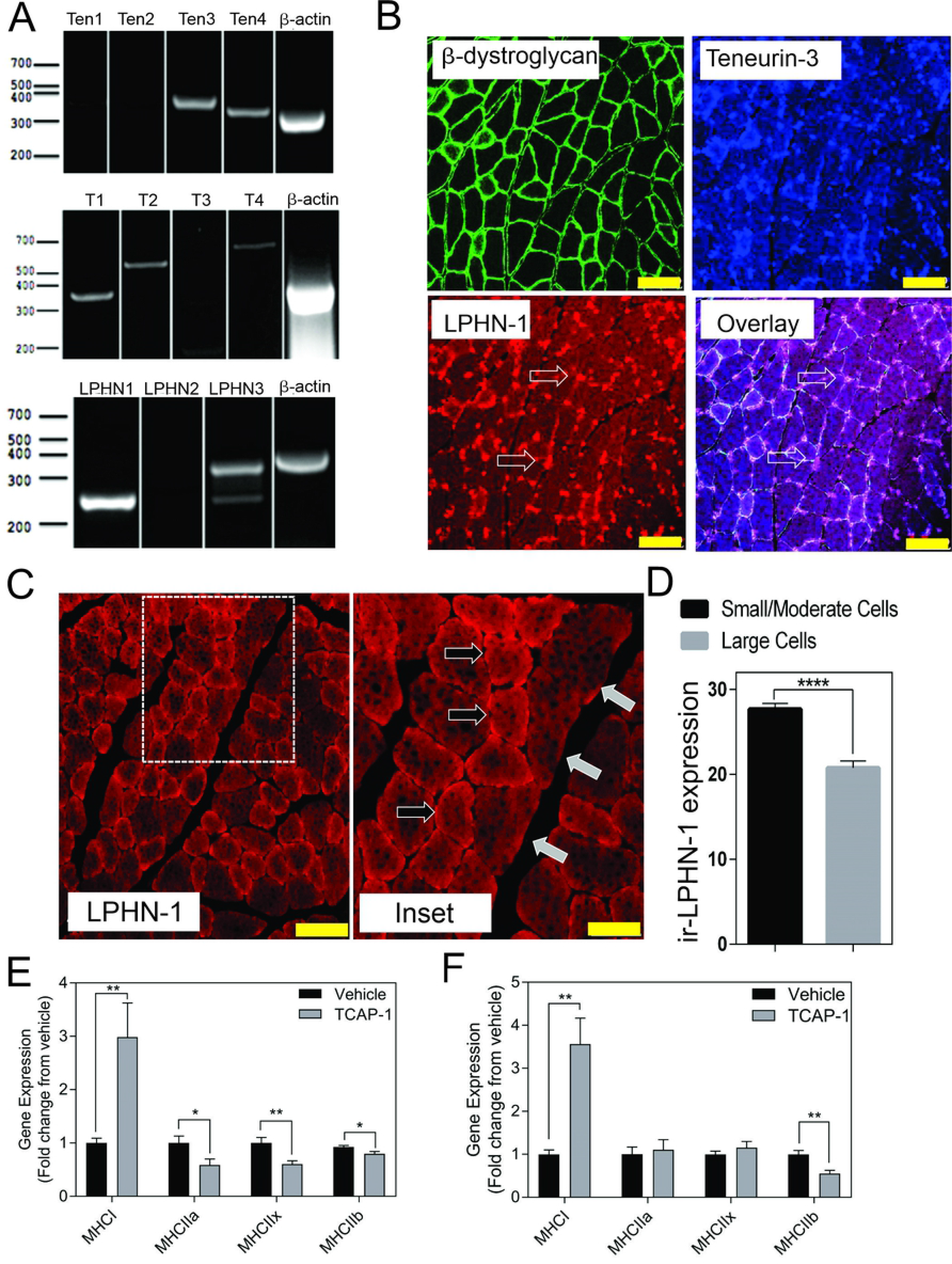
Expression of the teneurin/TCAP-LPHN immunoreactivity (ir) in rat skeletal muscle: **A**. PCR expression of teneurins, TCAP and LPHNs in rat TA muscle. **B.** Immunological expression of β-dystroglycan, teneurin and LPHN in rat skeletal muscle. Arrows indicate nodes of aggregation (Scale bar: 100 µM). **C.** Enhanced examination of ir-LPHN regions in TA muscle cells. Left panel scale bar indicates 100 µM, whereas the right panel scale bar indicates 50 µM. Black arrows indicate cells with high LPHN-1 labelling, whereas white arrows indicate cells of low LPHN-1 labelling. **D.** Quantification of ir-LPHN-1 as a function of muscle cell diameter (size) as shown in ‘C’ (Student’s t-Test p<0.0001; ****). **E.** Changes in fiber type in short-term TCAP-1 administration (t-test indicated for each pair). **F**. Changes in fiber-type over long term TCAP-1 administration (t-test indicated for each pair).

To corroborate these cDNA studies, immunohistochemical (IHC) studies were performed in rat TA muscle tissue. Initially, β-dystroglycan (DG) labelling was used to establish the sarcolemmic boundary of the cells, as previous studies indicated a relationship between DG and TCAP signaling [33]. Further, IHC co-localization labeling of Teneurin-1,-3 -and LPHN-1 was utilized to establish the cellular anatomical relationship between the teneurins and LPHNs. Immunoreactive (ir) teneurin-1 did not show a strong signal (data not shown), consistent with the RT-PCR data indicated above, however the ir-teneurin-3 showed a clear response indicating specific nodes of aggregation in the sarcolemma (Fig. 3B). Importantly, ir-LPHN-1 labelling of these tissues showed co-localization with the ir-DG along with ir-teneurin and ir-TCAP labelling consistent with the PCR studies indicated in Fig. 3A. The variation among the PCR and ir-teneurin -1 and -3 expression was expected due to affinity differences among the antibodies and primers (see discussion). Moreover, these studies establish a clear relationship between TCAP, teneurins and LPHNs in rat skeletal muscle (Fig. 3B). Morphological differences, with respect to cross-section diameter between the vehicle- and TCAP-1-treated animals could be established. Thus, TCAP-1 administration induced a 25% increase (p<0.001) between the number of small and intermediate cells relative to the untreated vehicle rats (Fig. 3C, D). Because small and intermediate fibers are typically oxidative muscle fibres, these observations suggested that TCAP could stimulate glucose uptake in skeletal muscle. To corroborate these findings, the expression of myosin heavy chain (MHC) was evaluated in the TA muscle of both short-term and long term TCAP-treated animals. In both cases, there was a significant (p<0.01) 3- to 3.5-fold increase in the expression of the MHCI fibres in the TCAP-1-treated animals compared to the non-treated vehicle, although significant (p<0.05) differences were also observed among MHCIIa, MHCIIx and MHCIIb expression among the treated and untreated animals (Fig. 3E,F).

Taken together, these studies indicated that TCAP-1 may increase glucose transport into skeletal muscle. Therefore, TCAP-1-induced glucose uptake into the hind-limb was measured directly by fPET. Thus, using ^18^F-deoxyglucose (FDG), a single dose of SC TCAP-1-treatment-induced FDG uptake in the hind-limb muscle by 2-fold (p<0.05) (Fig. 4.A,B) after 3d of treatment relative to vehicle treatment. These data corroborated our supposition that TCAP-1 acted, in part, to increase glucose importation into skeletal muscle. If this was the case, then this increase in glucose importation should increase skeletal muscle NADH production as a result of 2-glyceraldehyde-3-phosphate conversion to 2-1,3 diphosphoglycerate and secondarily through elements of the tri-carboxycyclic (TCA) acid cycle in the conversion to pyruvate. TCAP-1-treated muscle significant (p<0.05) increased NADH-staining compared to vehicle (Fig. 4C, D) supporting increased TCAP-1-mediated glucose transport.

**Figure 4.**
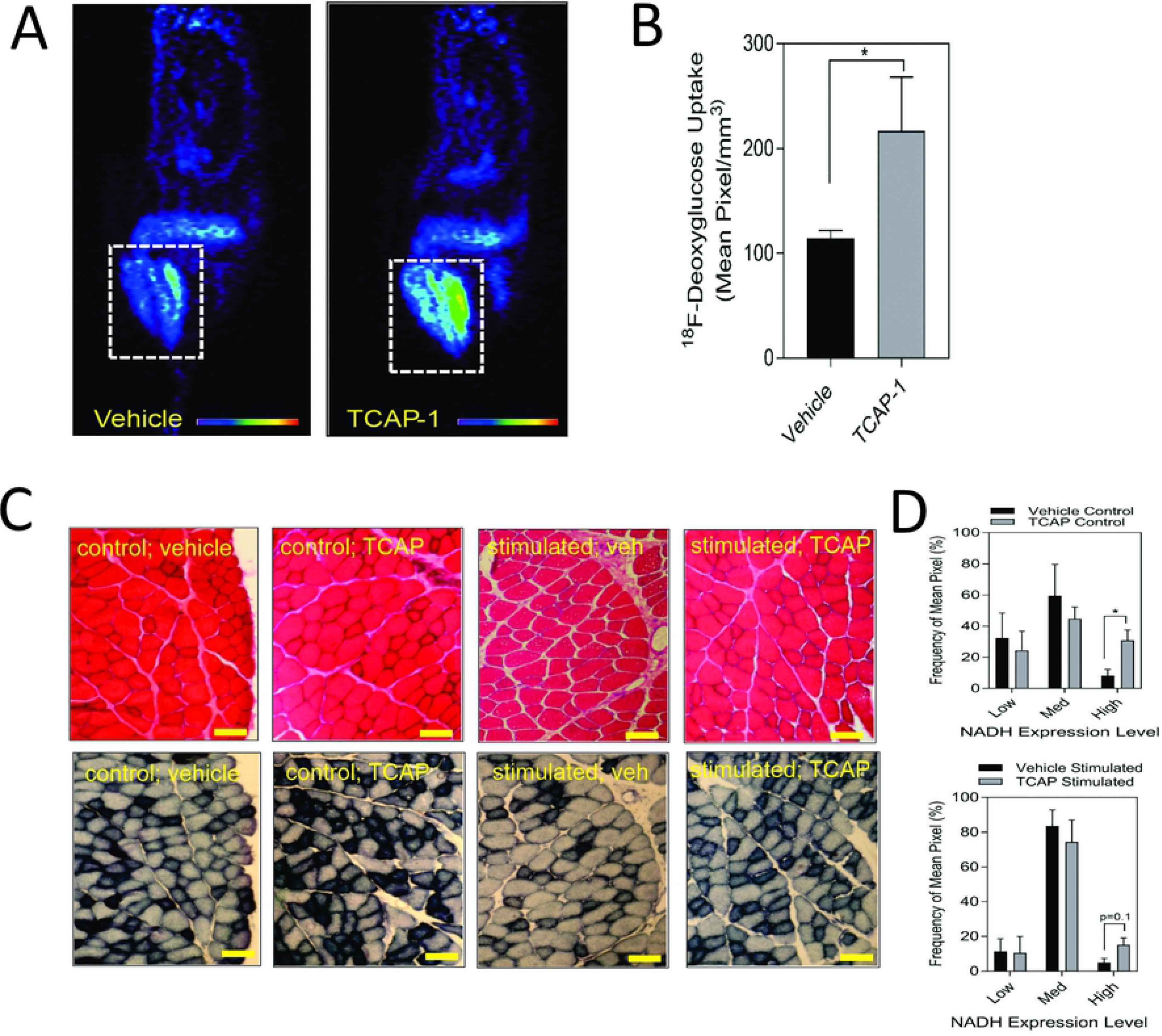
Glucose uptake and metabolism in rat hind-limb and TA muscle. **A.** Function positronic emission tomography (fPET) of rat hind limb showing increase of ^18^F-deoxyglucose (^18^F-DG) uptake after 3 days from TCAP-1 administration. **B.** Quantification of the ^18^F-DG uptake in hindlimb after 3 days. (n=5; student’s t-test; p<0.05). **C.** Stimulation of NADH in TA muscle after administration of TCAP-1. Above panel, (hematoxylin and eosin stain), bottom panel, NADH activity (shown as black regions). Scale bar indicates 100 µM. **D.** Quantification of the NADH-labelled cells shown in ‘C’. * p<0.05; **p<0.01; ***p<0.001; ****p<0.0001. Mean ± SEM indicated n=4.

Muscle performance is related to the amount of energetic substrates available, therefore, we examined the role of TCAP-1 on muscle activity *in vivo* in rats by determining the efficacy of TCAP-1-mediated contractility using electrical stimulation of the TA muscle. After a 5-d daily treatment of either vehicle or TCAP-1, following a 3-d washout period, muscle contractility was assessed. TCAP-1-treated animals showed improved muscle dynamics. TCAP-1-treated animals exhibited enhanced baseline contraction kinetics with respect to increased peak twitch force (p<0.05) (Fig. 5 A,B), slower contraction velocity (p<0.05) (Fig 5C), and potentially higher faster relaxation rates (Fig. 5D) compared to vehicle-treated animals. Following baseline measurements, a 6-m fatigue protocol was induced in the muscle where contractile kinetics were recorded at 0, 1, and 5m after the fatigue protocol. TCAP-1 enhanced recovery from the twitch stimulation (Fig. 5E-G). Although TCAP-1 did not influence peak twitch force (Fig. 5E), it significantly (p<0.05) maintained twitch max dx/dt (Fig. 5F) and 1/2RT (Fig. 5G) over the course of the fatigue protocol which was diminished in vehicle-treated animals. All data was normalized to muscle mass. The treatment did not affect muscle mass (Fig. 5H), tetanic force (Fig. 5I) or the fatigue force curve (Fig. 5J). Thus, TCAP-1 enhanced the efficiency of the existing muscle morphology, rather than increasing muscle mass, and maintained contraction cycling efficiency during fatigue. To assess the effects of a long-term (LT) treatment, rats were administered either vehicle or TCAP-1, for 3 months (1 injection/week). Two weeks post-treatment, the TCAP-1-treated animals elicited a comparable peak twitch force to vehicle-treated animals (Fig. 5K-J), however had significantly (p<0.05) slower contraction velocity and faster (p<0.05) relaxation rate (Fig. 5L,M).

**Figure 5.**
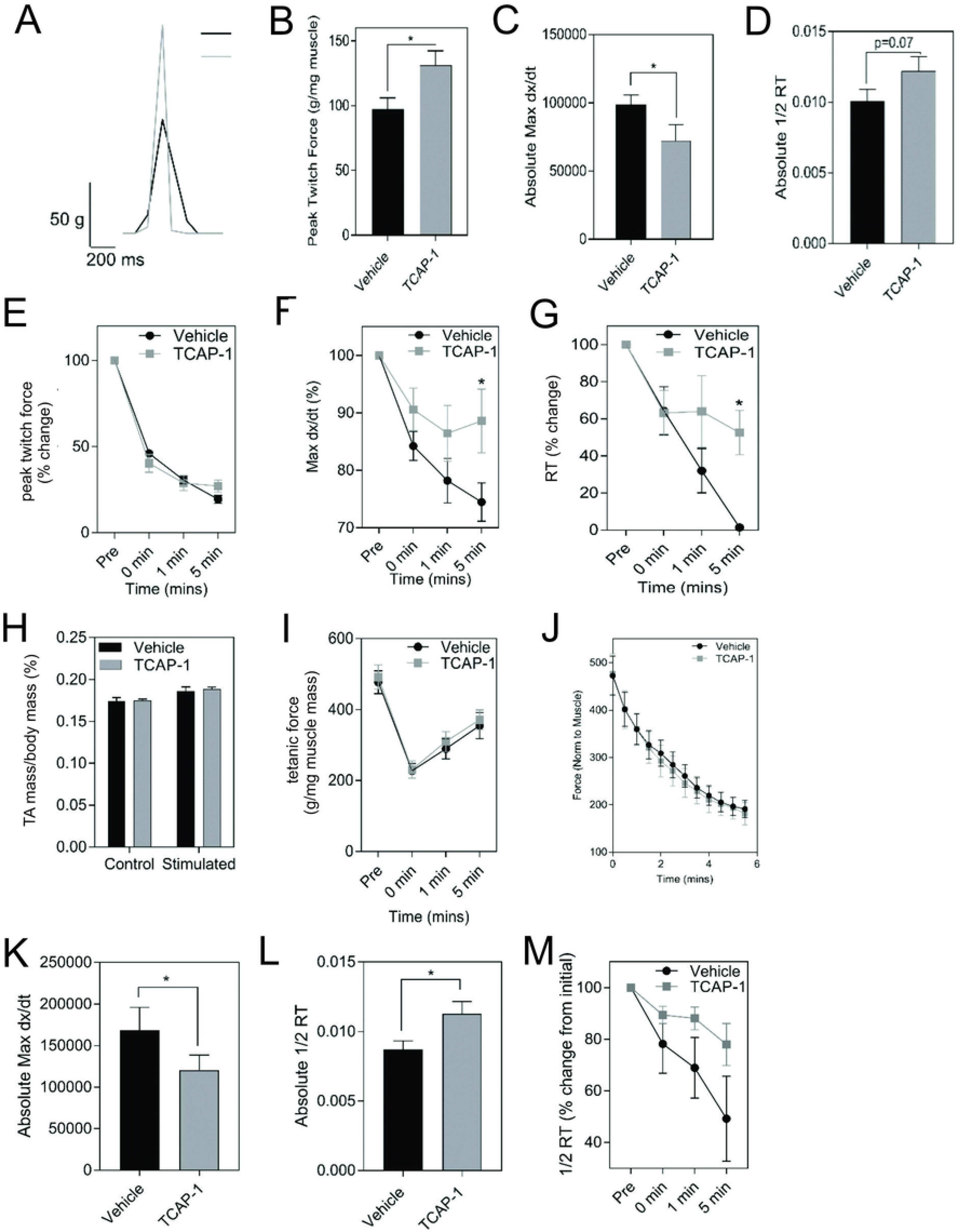
*In vivo* actions of TCAP-1 on rat TA muscle kinetics. **A-J**. Muscle twitch kinetics after animals treated with TCAP-1 once a day for 5 days before testing **A**. Representative twitch traces (black, vehicle; gray, TCAP-1). **B**. Baseline contraction kinetics (t-test). **C.** Contraction velocity (t-test). **D.** Relaxation rate (t-test). **E.** Peak twitch force (2-way ANOVA). **F.** Twitch max dx/dt (2-way ANOVA). **G.** 1/2RT analysis (2-way ANOVA). TCAP-1 treatment did not affect muscle weight (t-test) (**H**), tetanic force (t-test) (**I**) or fatigue force over time (**J**) (n=7-8). **K-M.** Long term treatment of TCAP-1 on rat hind-limb twitch kinetics. **K.** contraction max dx/dt. **L.** 1/2RT rate. **M.** Relaxation rate.

### In Vitro Mouse Cell Studies

The initial PCR screen of C2C12 cells indicated that, although only teneurin-3 was highly expressed (Fig.6A), all 4 TCAP transcripts could be discerned (Fig. 6B). In both undifferentiated C2C12 myoblasts, and 6-d myotubules, the transcripts for LPHN-1 and -3 were present (Fig. 6C). IHC expression of TCAP-1 showed a similar punctate expression in the cytosol of the C2C12 myoblasts (Fig. 6D) as we have previously shown for neurons [38] where FITC-labelled TCAP-1 was present at several sarcolemmic regions consistent with the expected expression of the receptor [32,33,38]. Moreover, because TCAP-1 regulates actin organization and polymerization in neurons [38], the C2C12 cells were treated with TCAP-1 and examined using the phalloidin stain to highlight actin fibers (Fig. 6E). This resulted in a major increase in actin polymerization in the TCAP-1-treated cells at both 30m (p<0.01) and 2d (p<0.001).

**Figure 6.**
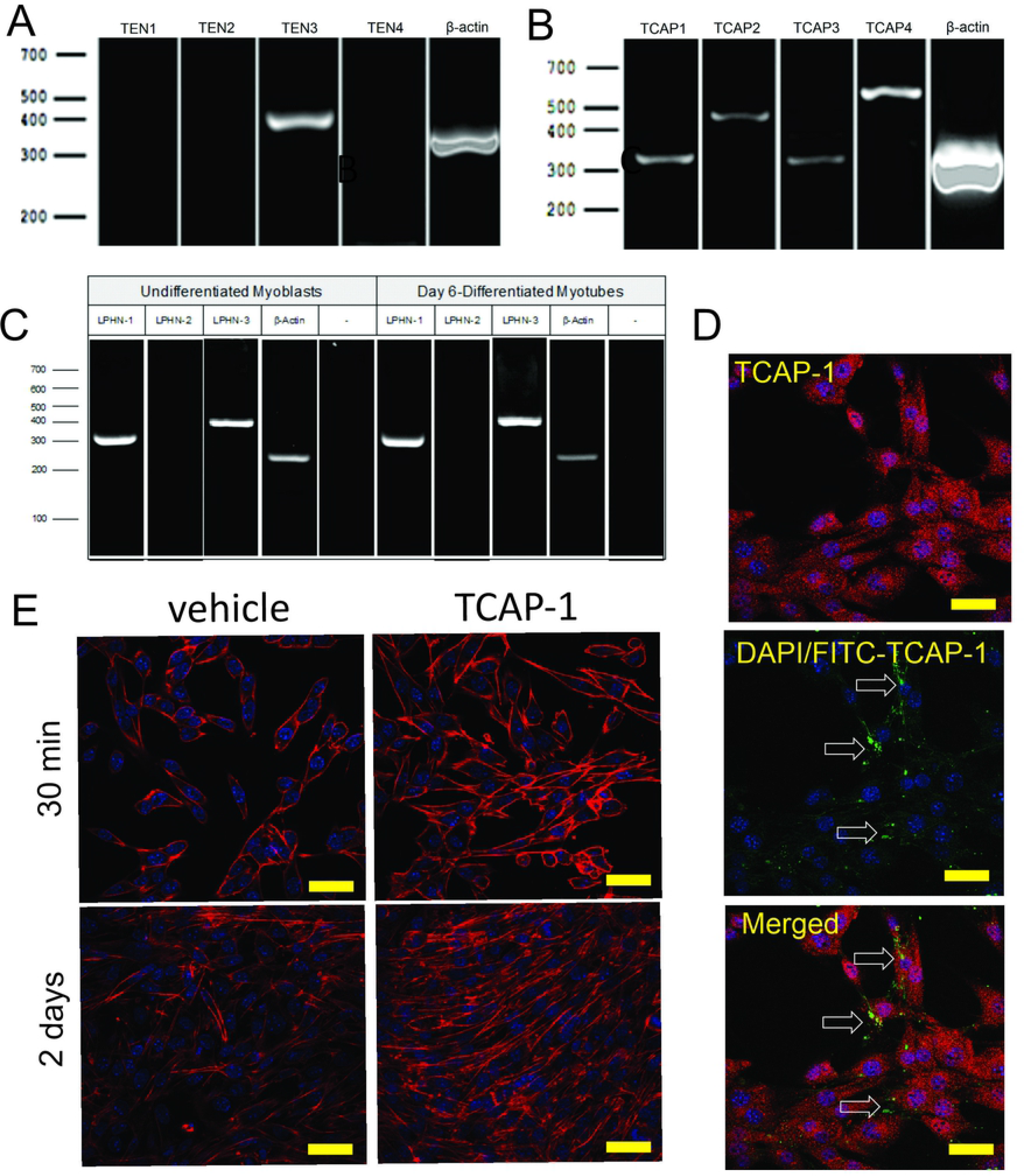
Expression of teneurin, TCAP and LPHN in C2C12 myoblasts. **A**. PCR-based teneurin expression; **B**. PCR-based TCAP expression; **C.** PCR-based LPHN expression. **D.** C2C12 cells labelled with TCAP-1 antisera showing the difference between the endogenous ir-TCAP and the presence of FITC-TCAP-1 localization. ir-TCAP-1 is indicated in red, whereas the DNA-associated DAPI labelling is indicated in blue. FITC-labeled TCAP-1 is shown in green. Arrows indicate regions of FITC-TCAP-1 uptake. Scale bars indicate 50 µM **E.** Actions of TCAP-1 on the proliferation of the C2C12 myoblasts when treated with TCAP-1 at 30 minutes and 2 days. Actin is indicated in red, whereas the nuclei are indicated in the DAPI-based blue. Scale bars indicate 100 µM.

Having established that TCAP-1 behaved in a similar manner as previously shown in neurons, the viability of the Ca^2+^ response in the differentiated myotubules was evaluated to determine their efficacy before proceeding to further studies. Initially, caffeine was used to determine the limits of the Ca^2+^ response in the differentiated myotubules relative to the TCAP-1 response (Fig. 7A-D). These studies indicated that the myotubules were active and viable, and with respect to the Ca^2+^ response, did not show an appreciable decrease in cytosolic Ca^2+^ concentrations (Fig. 7B), although it did attenuate the rate of cytosolic Ca^2+^ concentrations (p<0.01). Taken together, these studies indicated that the myotubules were viable with respect to our preparation, and that the attenuating TCAP-1 response indicated that additional regulating factors were likely present. Thus, given these observations, the direct action of TCAP-1 on Ca^2+^ flux in myotubules was examined (Fig. 7E-H). TCAP-1 increased Ca^2+^ concentrations by almost 4-fold relative to the control-treated cells (p<0.001).

**Figure 7.**
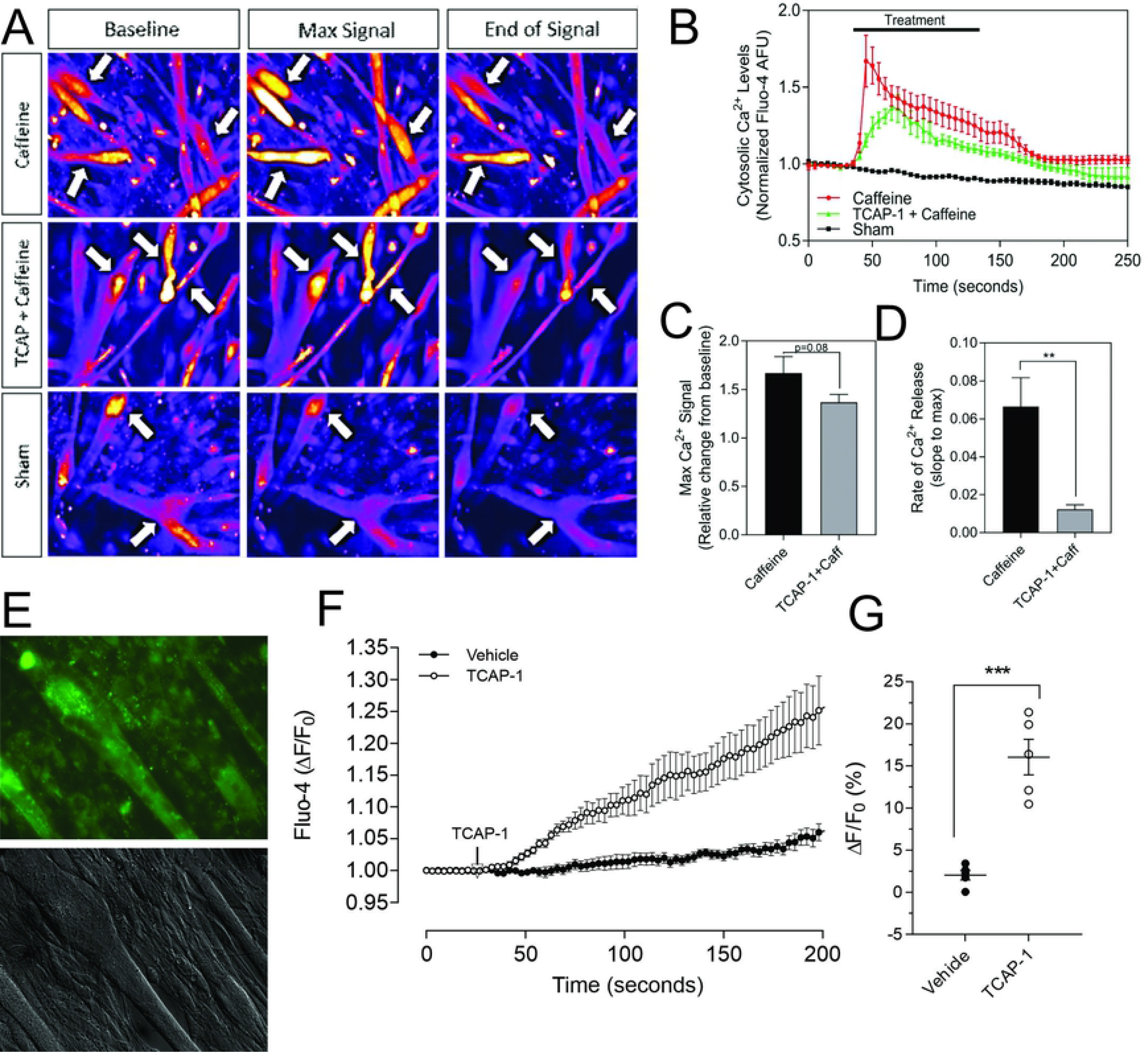
Caffeine- and TCAP-1-mediated Ca^2+^ response in differentiated C2C12 myocytes. **A.** Heat-map images showing the Ca^2+^ response induced by caffeine, TCAP-1 or vehicle. Mean and SEM is indicated**. B**. Dynamic concentration changes over the period of analysis shown in ‘A’. Mean ± SEM is indicated**. C.** Total concentration changes of the manipulations of the study period indicated in ‘A’. **D.** Rate of Ca^2+^ release between caffeine and TCAP-1. **E.** TCAP-1 mediated Ca^2+^ actions show normal morphology in cells. **F.** Rate of increase in Ca^2+^-associated fluorescence after administration of TCAP-1. **G.** Quantification of the change in TCAP-1 mediated intracellular Ca^2+^ concentrations indicated in ‘F’. Significance was determined by a Students t-test. (* p<0.05; **p<0.01; ***p<0.001; ****p<0.0001. Mean ± SEM indicated n=4).

In skeletal muscle, the predominant glucose transporter protein (GLUT) isoform is the insulin-sensitive GLUT4 protein. Using insulin as a control, a significant (p<0.001) increase in the expression of the ir-GLUT4 transporters was observed for both TCAP-1 and insulin treatments over 30m (Fig. 8 A,B) in C2C12 cells. To determine whether this GLUT4 response was dependent on the TCAP-1-mediated Ca^2+^ release, the IP3R antagonist, 2-APB, that abolishes the TCAP-1 Ca^2+^ response, was investigated. In the presence of the inhibitor, both TCAP-1 (p<0.01) and insulin (p<0.001) inhibited the ir-GLUT4 expression (Fig. 8A,C). This importation of glucose by TCAP-1 was further corroborated in C2C12 cells showing that TCAP-1 significantly (p<0.0001) induced ^3^H-2-deoxyglucose increase into the cytosol over 30m, similar to that of insulin (Fig. 8D). However, both peptides show distinct glucose-uptake profiles, where insulin induces a significant increase at 30m (p<0.001), 45m (p<0.001) and 60m (p<0.01), whereas TCAP-1 induces an increase at 30m (p<0.001) but was attenuated by 45m (p<0.01) and returns to baseline at 60m. The sc-TCAP-1 treatment, used separately as a negative peptide control in this study, showed no significant change from the saline vehicle.

**Figure 8:**
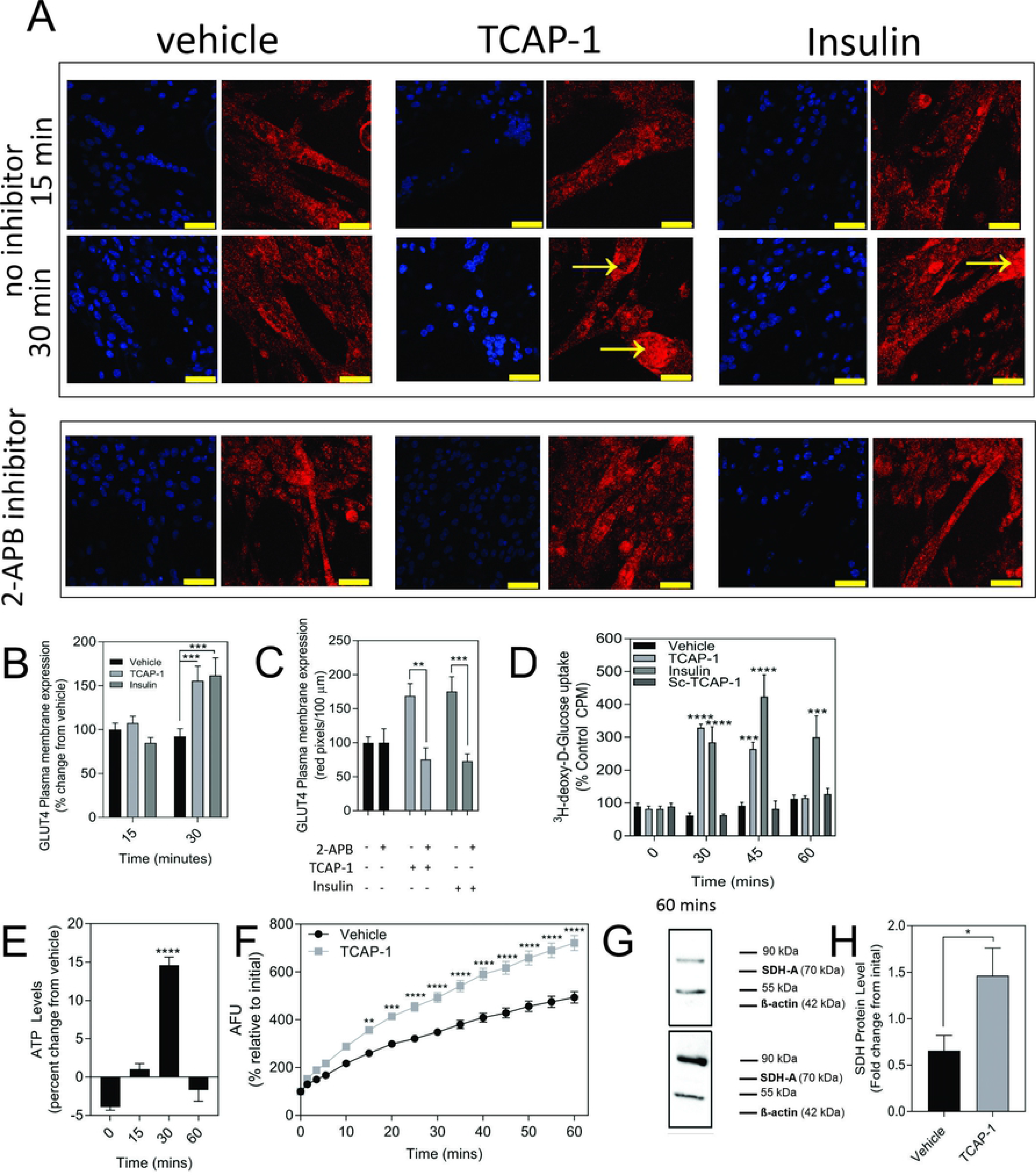
TCAP-mediated glucose metabolism in C2C12 cells. **A**. Regulation of ir-GLUT4 by TCAP-1 and insulin myotubules. Red indicates ir-GLUT4 whereas blue shows DAPI staining of the nuclei. Arrows indicated regions of high immunoreactivity. Scale bar=100 µM. **B**. Quantification of ir-GLUT4 expression over 30 minutes. **C**. Effect of the IP3R inhibitor, 2-APB on TCAP-1- and insulin-mediated ir-GLUT4 labelling. **D.** Uptake of ^3^H-2-deoxyglucose in C2C12 myoblasts by TCAP-1 and insulin. **E.** Changes in static ATP concentrations following treatment by TCAP-1. **F.** NADH production increase as determined by a resesourin assay following TCAP-1 treatment relative to the vehicle. **G**. Increased ir-succinate dehydrogenase expression after 1 hour following TCAP-1 treatment as determined by western blot. **H.** Quantification of the data indicated in ‘G’. Significance was determined by a t-test as indicated in C, F and H or one-way ANOVA shown in B D and E. (* p<0.05; **p<0.01; ***p<0.001; ****p<0.0001. Mean ± SEM indicated n=6).

Increased glucose importation increases ATP and NADH turnover in cells due to glycolytic and tricylic acid (TCA) cycle activity. Therefore, this was examined with respect to TCAP-1 treatment. As a result, both ATP (p<0.001) (Fig. 8E) and NADH (p<0.001) (Fig. 8F) turnover were significantly increased after 30m of TCAP-1 treatment, although NADH levels remained about 60% higher (p<0.001) than vehicle levels after 60m. Moreover, succinate dehydrogenase (SDH; a rate-limiting step of the TCA cycle) protein expression was increased by over 2-fold after TCAP-1 treatment corroborating the previous experiments of increased ATP and NADH production (p<0.05; Fig. 8G,H).

As we have previously established that the IP3-DAG pathway is important for TCAP-1-mediated intracellular Ca^2+^ flux in neurons, this pathway was examined in C2C12 cells. Relative to the vehicle, TCAP-1 induced a significant increase at 5 and 15 min (p<0.001; Fig. 9A) in intracellular DAG concentrations and a major increase between 1 and 15 min (p<0.0001; Fig. 9B) in IP3 concentrations. To confirm this action, TCAP-1-treated C2C12 cells were blocked with either the IP3R antagonist, 2-APB, or the phospholase C inhibitor, U73122. The 2-APB and U73122 treatment reduced TCAP-1-mediated Fluo-4 concentrations to about 30% (p<0.01) of their original values indicating that the IP3-DAG pathway plays an active role in increase of TCAP-1-mediated intracellular Ca^2+^ flux (Fig. 8C,D). Because this TCAP-1-mediated rise in intracellular Ca^2+^ concentrations can target the mitochondria [39], the mitochondrial Ca^2+^ dye, rhodamine 1-AM (Rhod-2) was utilized to determine the concentration of Ca^2+^ sequestration in mitochondria. There was a 5-fold increase (p<0.001) in Rhod-2-associated Ca^2+^ labelling over 200s (Fig. 9E,F). Related to this, Rhod-123, was used to determine the level of mitochondrial polarization. TCAP-1 treatment significantly decreased Rhod-123 fluorescence (p<0.001) relative to vehicle indicating depolarization of the mitochondria membrane (Fig. 9G, H).

**Figure 9.**
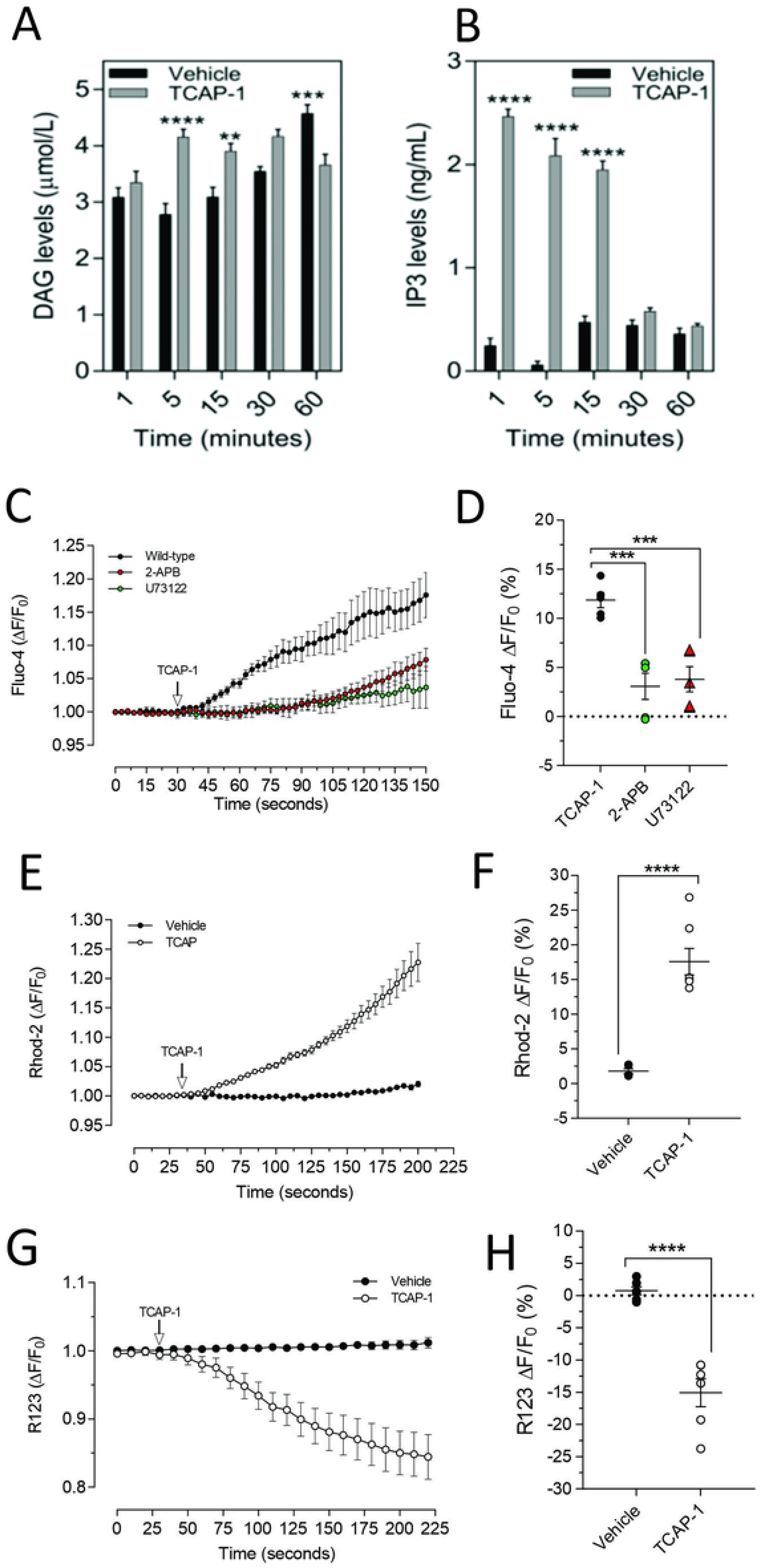
TCAP-1 mediated calcium regulation in C2C12 cells. **A.** TCAP-1 mediated increase in intracellular DAG concentrations (mean and SEM shown; n=6). **B.** TCAP-1 mediated increase in intracellular IP3 concentrations (mean and SEM shown; n=6). **C.** Increase in intracellular TCAP-1 mediated Ca^2+^ concentrations and inhibition by the IP3 receptor (2-APB) and phospholipase C (U73122) antagonists; **D.** Quantification of data shown in C (n=6). **E.** Uptake in Ca^2+^-mediated Rhod-2 into mitochondrial membranes. **F.** Quantification of data shown in E based on the change at 200 seconds. **G.** Decrease in Rhod-123 immunofluorescence in mitochondrial membranes as a result of TCAP-1 administration. This decrease in Rhod-123 indicates a decrease in mitochondrial membrane depolarization. **H**. Quantification of the data indicated in G based on the changes at 225 seconds. (* p<0.05; **p<0.01; ***p<0.001; ****p<0.0001.)

Although these studies are similar with previous investigations of teneurins, TCAPs and LPNHs in neurons, this is the first study to examine teneurin/TCAP and LPHN activity in skeletal muscle function. To determine whether the TCAP-1 activation was dependent upon the LPHN receptors, these genes were knocked-down (KD) using siRNA oligonucleotides, or knocked-out (KO) using CRISPR in the C2C12 cells. Using the C2C12 cells, the LPHN-1 and -3 expression was reduced using the siRNA oligonucleotides. The LPHN-1 receptor mRNA was reduced about an 80% (p<0.01) relative to the WT cells. Transfection with either the LPHN-1 siRNAs or the null vector (NT) did not significantly change mRNA expression relative to the WT control (Fig. 10A). Similarly, the LPHN-3 siRNA-associated oligonucleotides significantly (p<0.01) decreased its mRNA expression about 65% relative to the WT cells. There were no significant changes in mRNA expression of the LPHN-1 transcript in either the LPHN-1 KD or the NT cells (Fig. 10B). Despite the reduced expression of these receptors, cell morphology was normal (Fig. 10C). TCAP-1 increased cytosolic Ca^2+^ in cells transfected with the NT control, however, relative to the NT control, TCAP-1 did not increase Ca^2+^ in either the LPHN-1 and -3 siRNA-transfected cells, which showed a significant decrease (p<0.01 and p<0.001, respectively) in intracellular Ca^2+^ concentrations (Fig. 10D,E). However, because both LPHN-1 and -3 siRNA oligonucleotides unexpectedly reduced intracellular Ca^2+^ concentrations by similar amounts, we repeated this study by ablating the LPHN-1 and -3 receptors using CRISPR methods. The E5U7 target reduced LPHN-1 expression by about 90% (p<0.01) whereas, the E5D3 target reduced mRNA levels by about 95% (p<0.05) relative to the WT cells (Fig. 11A). A significant decrease (p<0.05) in LPHN-1 mRNA levels were present in both sets of transgenic cells (Fig. 11B). Importantly, cell morphology was normal in the transgenic cells (Fig.11C). CRISPR-based KOs of the LPHN-3 gene were unsuccessful after numerous attempts (data not shown; see discussion), hence studies were performed with the LPHN-1 KOs only. Similar to that achieved using the siRNA knock-down cells, both CRISPR-associated KO transgenic cells (E5U and E5D) reduced intracellular Ca^2+^ levels by about 60% (p<0.001) (Fig. 11 D,E). Taken together, both the siRNA- and CRISPR-associated methods to reduce mRNA expression indicate that the TCAP-1 associated intracellular Ca^2+^ flux was mediated primarily by its interactions on LPHN-1 and -3.

**Figure 10:**
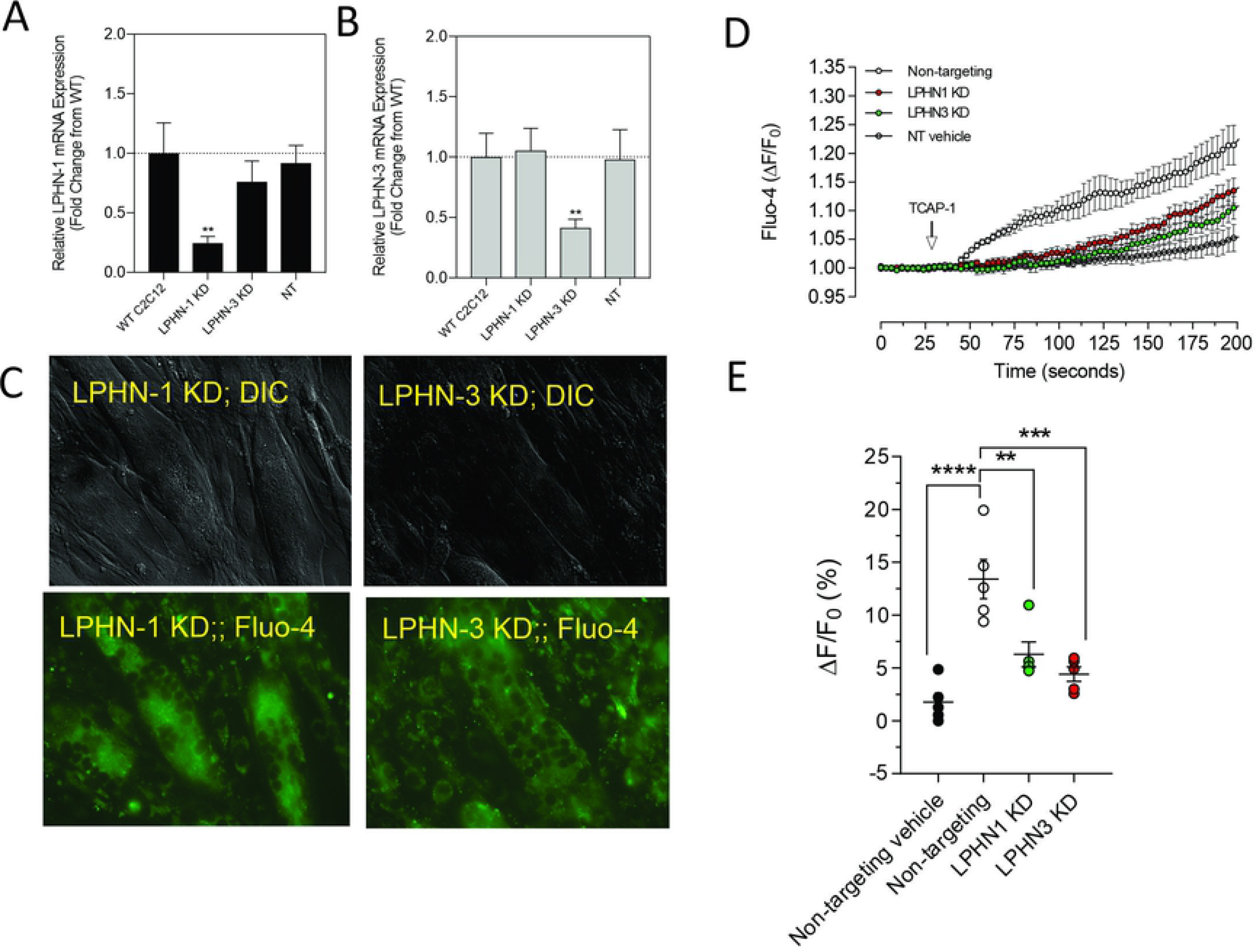
siRNA knockdown of LPHN-1 and -3 in C2C12 cells. **A.** Cells transfected with the LPHN-1-targetting siRNAs showed a significant reduction (p<0.01) in LPHN-1 mRNA expression. **B**. Cells transfected with the LPHN-3 targetting mRNA significantly (p<0.01) reduced LPHN-3 mRNA expression. C. Cells treated with either siRNAs showed normal morphology. **D**. Changes in Ca^2+^ accumulation in cells transfected with either LPHN-1 or -3 siRNA oligonucleotides. **E**. Quantification of the data shown in ‘D’ at 200 seconds.

**Figure 11:**
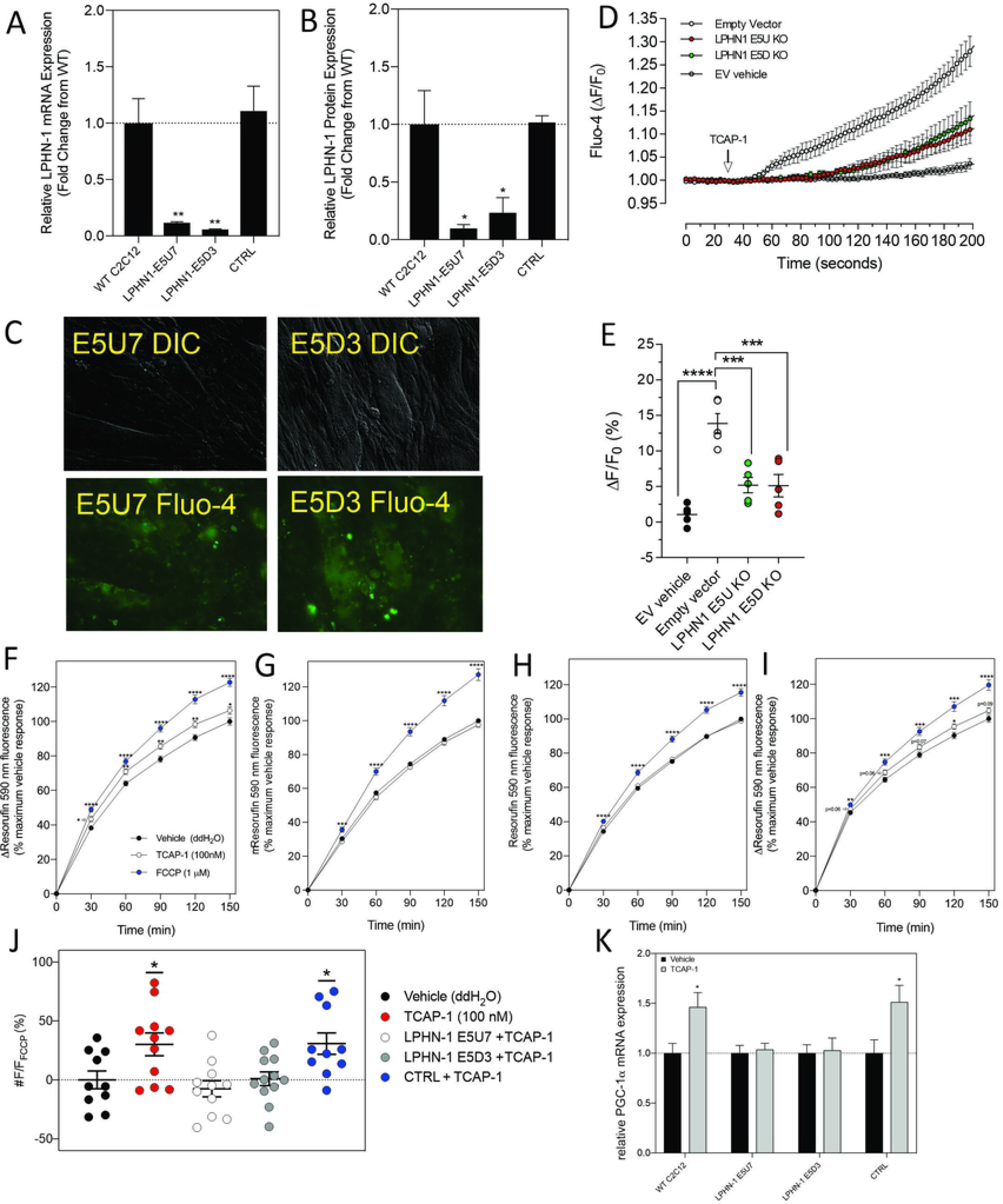
CRISPR-based knockouts of the LPHN-1 and -3 genes in C2C12 cells. The two clones (E5U7) (**A**) and (E5D3) (**B**) significantly reduced LPHN-1 expression relative to NT and WT cells. **C**. The morphology of the C2C12 cells were normal in both transfected cell lines. **D**. Changes in cytosolic Ca^2+^ accumulation in the various cell types. **E.** Quantification of the data shown in ‘D’ after 200 seconds. **F, G, H,I.** Changes in NADH production as determined by the resasurin assay **. J.** Quantification of the data shown in F-I, K. Reduction in PGA-1α mRNA expression by PCR in E5U7 and E5D3 clones. * p<0.05; **p < 0.01; ***p<0.001; ****p < 0.0001. Mean ± SEM indicated n=7-8.

To corroborate these findings with the previous observations that TCAP-1 could regulate energy metabolism in skeletal muscle cells, the action of TCAP-1 on NADH production via a resazurin assay was performed on the CRISPR LPHN KO cells (Fig 11F-I). From 30-150m, the TCAP-1-treated WT C2C12 cells showed a significant increase (p<0.01; p<0.05, respectively) in fluorescence compared to the vehicle (Fig. 11F). LPHN-1 E5U7 (Fig. 11G) and E5D3 (Fig. 11H). KOs did not show an increase after TCAP-1 treatment, whereas the NT control cells showed a significant (p<0.01 increase after 120m (Fig. 11I). FCCP treatment, indicating cell viability induced significant increases (p<0.001; p<0.0001) across all cell types (Fig. 11 F,G,H,I).

Given that the *in vivo* studies showed that TCAP-1 can modulate MHCI and that TCAP-1 may also affect other hormones and signalling factors, it was unclear if the purported fibre changes observed in TA muscle was a direct result of TCAP-1. Therefore, because PGC-1α is a transcription factor that up-regulates MHCI expression [46], the TCAP-1 actions on PGC-1α was measured by qRT-PCR. Following the TCAP-1 treatment (100nM), neither the E5U7 nor the E5D3 cells showed any significant increases, whereas, both the WT cells and NT controls showed about a 50% increase (p<0.05) in PGC-1α expression (Fig. 11K). This indicated that TCAP-1 has the potential to directly influence MCH fibre expression at the transcriptional level.

## Discussion

This study describes a novel mechanism underlying skeletal muscle physiology. This investigation is the first to show a functional relationship between teneurins and latrophilins (LPHN) with respect to skeletal muscle function in mammals using rodent models. We have previously established that there is a functional peptide on the distal tip of the teneurin extracellular region which we have termed ‘teneurin C-terminal associated peptide’ (TCAP) and is highly active in the CNS. Now, our data indicates that TCAP-1 affects skeletal muscle strength and fatigue, *in vivo,* via a glucose-associated and, likely, an aerobic mitochondrial-based mechanism. This mechanism is consistent with our previous findings in neurons and the CNS. Moreover, TCAP-1 interacts with the putative teneurin receptors, LPHN-1 and -3 to activate the PLC-IP3-DAG pathway to regulate intracellular Ca^2+^ flux that ultimately regulates glucose importation and mitochondrial activity. The hypotheses developed from the *in vivo* studies were subsequently tested *in vitro* utilizing the rat skeletal cell line, C2C12, to establish a cellular model upon which to base the *in vivo* actions of TCAP-1 with respect to skeletal muscle dynamics.

A critical aspect of this study was the utilization of TCAP-1 as a peptide analogue of the distal C-terminal region of teneurins. The genomic structure of the TCAP region of teneurins indicated that it possessed a potentially cleavable peptide [22,24]. A synthetic version of TCAP-1 was developed by replacing the N-terminal glutamine with pyroglutamyl acid and amidating the C-terminal residue [24]. The resultant synthetic peptide was highly efficacious at regulating neural function, behaviour and reproductive physiology in rodents [24,31,33,36-38] indicating that TCAP-1, itself, possessed independent biological functions. Thus, rat/mouse TCAP-1 was utilized in this study based on our previous work with this peptide. As indicated in Fig.1, the primary structure of rat and mouse TCAP-1 is identical, and consequently, the same peptide was utilized for all *in vivo* and *in vitro* experiments. However, it is important to point out that, although there are four paralogous forms of teneurins and TCAP in vertebrates, albiet, with significant primary structure conservation among them, our study has utilized only TCAP-1. TCAP-1 was utilized, therefore, as a proxy for all TCAPs present in the organism or tissue, to determine the potential to regulate skeletal muscle function. Based on our previous studies of this peptide and its level of primary structure conservation, evidence indicates that our supposition is valid.

Elucidation of the teneurin/TCAP-LPHN network is complex. To date, there are 4 teneurins found in vertebrates, each of which possesses a TCAP at its extracellular tip [4,17,18,23,24,26]. Moreover, the 3 latrophilin paralogues (LPHN1-3) have been identified in the vertebrate genome [29,47,48]. A clear stoichiometry among teneurins and the LPHNs has been only partially resolved. Both ligand and receptor proteins possess multiple domains that interact with a variety of peripheral ligands in the extracellular matrix, the membrane and intracellularly as well [10,13,14,21,22]. TCAP, as an amphiphilic peptide, that may be cleaved [14,22], or expressed separately [33,34] is a ‘wild-card’ in this perplexity of molecular interactions. Although TCAP is clearly bioactive with respect to cytoskeletal reorganization [32,33,37,38], glucose regulation [3,39], signal transduction [24,38,39,49], metabolism [24,32,34,39] and stress-associated behaviour [36, 50-54], these studies have focused at the neurological level. Despite this emphasis, few studies [55] regarding the role of teneurins/TCAP and LPHN on skeletal muscle physiology have been reported.

Our goal in examining the role of TCAP-1 and LPHN-1 and -3 was not intended to establish a specific molecular interaction, per se, but rather to show that the teneurin/TCAP-LPHN system plays an important role in skeletal muscle metabolism. Consequently, our investigation that paralogues of teneurins, TCAPs and LPHNs were present both in hind-limb skeletal muscle preparations and in C2C12 cells provided the basis of this molecular system with respect to skeletal muscle physiology. Because TCAP-1 co-localization to cell membranes was associated with the dystroglycan (DG) complex [38], we used DG antibodies to delineate the sarcolemma. In this study, the interaction of the teneurin-3 and LPHN-1 in the TA muscle occurred at specific nodes in the sarcolemma rather than being spread throughout all regions of DG labelling (see Fig. 3B). Our data indicates that teneurin-3 is the dominant teneurin in skeletal muscle tissue. However, the antisera available for the teneurins limited us to visualize histochemically only teneurin-3-like epitopes, although PCR expression indicated that the teneurin-3 transcript was expressed. All 4 TCAPs were expressed by PCR although TCAP-3 showed low expression. In contrast, expression of these transcripts in C2C12 cells indicated high expression of teneurin-3 along with all 4 TCAPs (see Fig. 6A,B). This is the first time we have observed high TCAP expression without the corresponding teneurin expression using PCR (Lovejoy, unpublished observations), although we have only focused on neurons previously. We acknowledge that it can be difficult to reconcile the expression patterns of teneurins using the antibodies available at the time, however, it is important that both elements are present. These data may indicate a fundamental difference among teneurin and TCAP expression in skeletal cells relative to neurons.

Our data indicated that TCAP-1, like in neurons [39], regulates glucose-mediated aerobic-based energy metabolism in skeletal muscle. Several observations support this hypothesis. First, *in vivo*, ir-LHPN primarily labeled small and intermediate muscle fibres in the rat TCA muscle (see Fig. 3C,D), that are typically associated with aerobic action. Second, with respect to MHC proteins, TCAP-1 showed the greatest increase in MCHI expression in both short-term and long-term TCAP-1 treatment of rats. Muscle kinetics are dependent, in part, on the relative proportion of muscle fibers types. Although the TA muscle consists of 95% type-II muscle (fast-twitch glycolytic muscle fiber type), TCAP-1 imparts traits of type-I fibers (slow-twitch and oxidative) which possess greater MHC expression. Importantly, both short-term (p<0.01) and long-term TCAP-1 (p<0.01) administration (Fig.3 E,F) increased in the MCHI fibre expression, although in short-term TCAP-1-treated animals, MHCII expression was specifically decreased in MHCIIa, (p<0.05); MHCIIx (p<0.01) and MHCIIb (p<0.05) mRNA expression, where only MHCIIB showed a decrease (p<0.01) in the long-term treated rats. These expressional changes of MHC transcription were corroborated by the expression of PGC-1α, a critical transcriptional co-factor that regulates the mitochondrial actions of myosin chain transcription. Using the C2C12 cells as a model, TCAP-1 increased the transcription of PGC-1α and was inhibited (p<0.05) by CRISPR-mediated KOs of the LPHN-1 gene. Thus the TCAP-1-mediated Ca^2+^ surge may activate CaMKIV and CaN transcriptional regulators to promote the transcription of PGC-1α. Overall, this indicates that TCAP-1 is increasing slow-twitch gene expression, consistent with the *in vivo* contractile kinetics observed in Fig 5. Third, rats treated with TCAP-1 enhanced baseline contractile kinetics under basal and fatigue conditions using both short- and long-term TCAP-1 administration. Fourth, a single dose of TCAP-1 increased uptake of ^18^F-2-deoxyglucose in rat hind-limb regions as determined by fPET analysis. Fifth, because Ca^2+^ and ATP are required for proper muscle contraction initiation and relaxation, TCAP-1 significantly increased GLUT4 expression (Fig.8A), increased ^3^H-2-deoxyglucose uptake (Fig. 8D), ATP (Fig. 8E) and NADH (Fig. 8F) concentrations and increased the protein expression of SDH in the TCA cycle (Fig. 8G,H) using the C2C12 myoblasts and myotubules. Taken together, these studies support the hypothesis that TCAP-1 regulates glucose uptake and metabolism in skeletal muscle cells in a similar manner previously described in neurons [39].

Glucose is the main energy nutrient for skeletal muscle function, thus, up-regulation of glucose metabolism could impact skeletal muscle activity. Therefore, the influence of TCAP-1 on these contractile kinetic parameters indicates that TCAP-1 modulates Ca^2+^ levels to enhance SR-sarcomere coupling. During fatigue, cytosolic Ca^2+^ levels accumulate due to inefficient SR-sarcomere coupling, thereby reducing muscle function as was observed in contractile kinetic parameters such as peak twitch force and 1/2RT. TCAP-1 treatment significantly increased both parameters, indicating a clear role in Ca^2+^ modulation. Moreover, these results also suggest that TCAP-1 increases ATP production rate to meet the energetic demands of the muscle during fatigue. After the contraction, Ca^2+^ is cleared from the cytosol and re-uptaken into the SR via the associated Ca^2+^ -ATPase (SERCA) pumps. SERCA pumps are high energy-consuming channels that account for 20-50% of the energy turnover in a single contraction cycle [43,56]. Prolonged stimulation results in the rapid depletion of ATP, thus reducing SERCA activity, ultimately leading to the accumulation of cytosolic Ca^2+^. However, because TCAP-1 increases glucose uptake into the muscle, it provides additional substrates for energy metabolism which could maintain SERCA activity during fatigue. This is corroborated by the finding that TCAP-1 had significantly faster 1/2RT during fatigue compared to vehicle treatment and by the increase in the type-1 muscle fiber associated gene transcription that occurred in both short-term and long-term actions of TCAP-1.

From these studies, it is clear that TCAP-1 targets the mitochondria as it has potent actions upon Ca^2+^ modulation and glucose signaling. As TCAP-1 increases Ca^2+^ uptake into the mitochondria, likely from shuttling Ca^2+^ from the SR, this stimulates enzymes in the TCA cycle, specifically glycerol phosphate dehydrogenase, pyruvate dehydrogenase phosphatase, isocitrate dehydrogenase and oxoglutarate dehydrogenase [57,58]. This activates mitochondrial respiration via the ETC and leads to increased energetic output, as seen by increases in ATP, NADH and SDH. Thus, this may explain enhanced metabolism and function results under TCAP-1 treatment. Previous studies in neurons have established that the IP3-DAG pathway is activated in response to TCAP-1. In this study, we showed that a similar situation occurs in C2C12 cells and, likely, in skeletal muscle. TCAP-1 treatment of C2C12 cells increase intracellular Ca^2+^ flux that can be blocked using IP3 receptor (2-APB) and phospholipase C (U73122) inhibitors. Moreover, this increase in intracellular Ca^2+^ is likely responsible for the depolarization of the mitochondrial membranes. This work supports a previous report that SR-associated Ca^2+^ release was directed toward the mitochondria in skeletal muscle [59].

These studies are consistent with previous observations in immortalized mouse neurons [39] and in zebrafish [34]. Although these previous studies indicate a relationship with TCAP-1 and LPHN action on Ca^2+^ flux involving the IP3-DAG pathway [39,60] other studies indicate that the LPHN-mediated AMP-PKA pathway may also be activated by teneurins and TCAP [61,62]. For example, TCAP-1 and -3 can increase cAMP levels in immortalized mouse neurons [24,37,49]. Further, studies of vertebrate teneurins have also implicated activation of the PKA-cAMP cascade [31]. However, our goal in this study was to establish a mechanism by which TCAP-1 can regulate energy metabolism in C2C12 cells, and for this reason we have focused on a Ca^2+^-associated mechanism, as it aligns with previous studies. However, we acknowledge that given the complexity of teneurin-LPHN actions, other signal transduction systems such as ERK-MEK [38] are also likely required for the full set of teneurin- and TCAP-mediated LPHN actions on cells. Ancient-evolving peptide-protein systems will likely impinge on more than one intracellular signal cascade events because they evolved before many of the later intracellular signaling transducing pathways [31].

The mitochondria are ultimately responsible for supplying aerobic-based energy requirements to eukaryotic cells. We have previously shown the relationship of TCAP-1 mediated energy production and the mitochondria in the mouse neurons [39] and in zebrafish metabolism [34], however this was the first study to establish the link between the teneurin/TCAP-LPHN system and mitochondria in skeletal muscle cells. Although we have not studied mitochondria respiration directly in this study, previously we showed that TCAP-3-treated zebrafish [34] increased both basal and respiratory reserve capacity. Although total mitochondrial respiration is linked to proton leak and ATP-linked respiration [63], no TCAP-3-associated actions on the latter could be detected, but proton leak was increased in these studies.

The role of the teneurins, TCAP and LPHNs, together, has previously not been examined in skeletal muscle. Therefore, we utilized both siRNA- and CRISPR-based methods to determine if the reduced activity of the LPHN-1 and -3 receptors would attenuate TCAP-1-mediated intracellular actions. Both CRISPR-based KOs of LPHN-1 significantly reduced the TCAP-1 associated increase in intracellular Ca^2+^. We were unsuccessful to create a LPHN-3 KO however. The length of the LPHN-3 gene in mice is about 10 times that of the LPHN-1 gene due to much longer intronic sequences. Thus, this extended sequence may have played a role in the lack of viability of the CRISPR-associated LPHN-3 oligonucleotides. Moreover, TCAP-1-mediated intracellular Ca^2+^ was established using pharmacological antagonists of the PLC-IP3-IP3R pathway of the SR. This rise in intracellular Ca^2+^ led to increased cellular glucose, concomitant with increases in ATP and NADH production. Indeed, the TCAP-1-mediated increase in intracellular Ca^2+^ corroborated with mitochondrial membrane hyperpolarization and increased succinate dehydrogenase activity indicating that TCAP-1 also acted to increase mitochondrial activity. However, the siRNA oligonucleotides did inhibit the actions of both receptors as indicated by the significant reduction in the TCAP-1-mediated cytosolic Ca^2+^ response. siRNA KDs of the LPHN-1 and -3 has been successfully used in the past using the mouse pancreatic β-cell line, MIN6 [64] where the authors showed that LPHN-3 KDs reduced insulin secretion by reducing cAMP levels via a Gi-mediated pathway. Although we have not examined the direct role of TCAP-1 on pancreatic insulin release, this study is consistent with our supposition that TCAP, itself, is associated with glucose regulation *in vivo* [3,39]. In our current study, however, our experiments showed that ablation of LPHN-1 or -3 reduced TCAP-1-mediated Ca^2+^ concentrations toward baseline levels. These results surprised us. We expected that the KO or KD or either receptor would render only a partial suppression of the Ca^2+^ response. Because this was not the case, one possibility is that there is an interaction among the LPHN isoforms specifically, or their combined actions with the teneurins.

Teneurin and LPHN interaction is complex where the stoichiometry among the 4 teneurin and 3 LPHN paralogues has not been ascertained. Although studies using vertebrate models establish clear evidence of teneurin-LPHN interaction [14,22,28,31], the specific correspondence of any teneurin with any LPHN as cognitive pairs has yet to be established. The teneurins are multifunctional transmembrane proteins that have TCAP at their distal extracellular tip. Even less is understood regarding the promiscuity of the LPHNs with respect to TCAP interactions. Previously, Silva and his associates [31] showed that the teneurin-2 region possessing the TCAP unit was required for full binding to the LPHN-1 (Lasso), and Husic and her colleagues [32] showed that the transgenic expression of teneurin-1 TCAP co-precipitated with the transgenic over-expressed hormone binding domain (HBD) of LHPN-1 and, moreover, modulated the cell adhesion characteristics of HEK293 cells overexpressed with the LPHN-1 mRNA. In this current study, we established that both LPHN-1 and teneurin-3 transcripts were present and are co-localized in the sarcolemma (Fig.3). These data corroborate with the PCR expression data indicating that these mRNA transcripts show the highest expression, but are not meant to suggest that this is indicative of cognitive ligand and receptor pairs, per se.

There is evidence that LPHN paralogues interact with each other. In our LPHN-1 and -3 attenuation studies, the reduction of one receptor inhibited the TCAP-1 mediated Ca^2+^ actions of the other receptor. It is possible that there is a minor Ca^2+^ response mediated by the non-target LPHN, but too low to detect with our assay conditions, although this seems unlikely. Homo-and heterophilic oligomerization is a characteristic of the GPCRs, but this has not been well-studied in Adhesion GPCR family members [65,66]. In LPHN-1, the C-terminal fragment and N-terminal fragment is cleaved *in vivo* and re-associates with the α-latrotoxin (αLTX) mutant ligand LTX^N4C^ [67], although this phenomenon has not been studied in detail in other LPHN paralogues despite the high degree of conservation among these domains. Further, studies using the ‘*stachel*’ peptide have provided additional insight into potential LPHN paralogue interactions with each other [64,68,69]. The *stachel* peptide represents the N-terminus of the C-terminal fragment following cleavage of the conserved GPCR-autoproteolysis (GAIN) region [70]. The recombinant expressed *stachel* peptide induces diverse G-protein associations across LPHN paralogues, whereas αLTX favours G11 signalling via LPHN-1 [60,71,72]. We and others have previously shown that the TCAP amino acid sequence resembles that of Secretin GPCR family ligands and αLTX [26], thus we posit that TCAP may represent the endogenous ligand that αLTX co-evolved with to become a toxin.

One of the questions we did not address in this study was the trigger that stimulates TCAP to regulate skeletal muscle physiology. Our hypothesis, at this time, is that TCAP is liberated locally in the sarcolemma either by a direct cleavage of the teneurin [14,22,23] or by independent mRNA transcription, translation and release from skeletal muscle or other local tissues [33,34]. We have shown in the past that TCAP-1 can increase teneurin transcription in immortalized neurons [24], but we have yet to establish this *in vivo*. Although TCAP is highly expressed in the brain, our studies showing its presence in as a circulating hormone in serum has been equivocal (Lovejoy, unpublished observations). As this is a critical aspect of TCAP action, this is a goal for upcoming studies.

In summary, our data in this study indicate that TCAP-1 regulates energy metabolism in skeletal muscle via an insulin-independent mechanism, and by doing so, modulates contractile kinetics, via Ca^2+^ dynamics and ATP production. Together, these data describe a previously unknown mechanism to regulate skeletal muscle dynamics. These data provide the foundation for a proposed mechanism of TCAP-1 action in skeletal muscle. TCAP-1 interacts with LPHN-1 and -3 to stimulate the activation of G-protein-coupled PLC leading to the increased conversion of PIP3 into IP3 and DAG. Increased IP3 levels stimulates the IP3R on the SR, opening Ca^2+^ channels to increase cytosolic Ca^2+^ levels. Cytosolic Ca^2+^ is imported into the mitochondria likely via the Ca^2+^ uniporter (MCU) which stimulates the TCA cycle and electron transport chain (ETC). Enhanced ETC activity results in increased proton extrusion from the mitochondrial matrix and hyperpolarization of mitochondrial membrane potential. This ultimately results in increased ATP and NADH production, as well as increased SDH-ATP levels. Ca^2+^ is subsequently pumped out of the mitochondria likely via Na^+^/Ca^2+^ exchangers (NCX), thus restoring homeostatic levels of Ca^2+^. Moreover, we showed that the TCAP-1 increased cellular energy availability by increased glucose importation into cells likely due to increased GLUT4 expression. The TCAP-1 mediated mechanism is likely due to its interactions with LPHNs -1 and -3.

## Acknowledgments

We thank the Canadian Natural Sciences and Engineering Research Council (NSERC) for Discovery Grants to Profs. D.A. Lovejoy, M. Locke and L. Buck and for NSERC Post-Graduate Scholarships to Dr. Andrea Reid, Yani Chen, Mei Xu and Mia Husic and an Ontario Graduate Scholarship to Thomas Dodsworth. We also thank Protagenic Therapeutics Inc. for operational funding to Prof. D.A. Lovejoy

## Author Contributions

Dr. Andrea Reid performed most of the experiments in this study, and wrote the initial draft of this paper. Dr. David Hogg developed many of the calcium-associated experiments and supervised the calcium studies and analyses, Dr. D. Barsyte and T. Dodsworth developed the strategy, preparation and analyses of the siRNA and CRISPR cell lines. Prof. P. Biga and R. Reid provided the guidance on the role of TCAP with respect to mitochondrial respirations and NADH assays, Dr. A. Slee provided critical information on TCAP-1 kinetics. Prof. L. Buck provided the guidance for the studies on energy metabolism in cells, and Prof. M. Locke developed and supervised the *in vivo* studies on muscle metabolism. Mei Xu designed and performed the initial experiments to determine the energy actions of TCAP in cells, Yani Chen developed the methods and analysed the uptake of glucose in muscle cells in *in vivo* and *in vitro*. Mia Husic established some of the methods for the cell culture of the transgenic cells and Prof. D. Lovejoy oversaw the entire research program and analyses, and prepared the final drafts of the manuscript.

## Notes

### Competing Interest Statement

The authors have declared no competing interest.

